# Identification of an H-Ras nanocluster disrupting peptide

**DOI:** 10.1101/2023.09.07.556635

**Authors:** Ganesh babu Manoharan, Candy Laura Steffen, Karolina Pavic, Alejandro Yeste-Vázquez, Matias Knuuttila, Neha Arora, Yong Zhou, Harri Härmä, Anthoula Gaigneaux, Tom N. Grossmann, Daniel Kwaku Abankwa

**Affiliations:** Cancer Cell Biology and Drug Discovery group, Department of Life Sciences and Medicine, University of Luxembourg, 4362 Esch-sur-Alzette, Luxembourg; Department of Chemistry and Pharmaceutical Sciences, VU University Amsterdam, Amsterdam, The Netherlands; Amsterdam Institute of Molecular and Life Sciences (AIMMS), VU University Amsterdam, Amsterdam, The Netherlands; Turku Bioscience Centre, University of Turku and Åbo Akademi University, 20520 Turku, Finland; Department of Integrative Biology and Pharmacology, McGovern Medical School, UT Health, Houston, TX 77030, USA; Chemistry of Drug Development, Department of Chemistry, University of Turku, 20500 Turku, Finland; Bioinformatics Core, Department of Life Sciences and Medicine, University of Luxembourg, 4367 Esch-sur-Alzette, Luxembourg

## Abstract

The Ras-MAPK pathway is critical to regulate cell proliferation and differentiation. Its dysregulation is implicated in the onset and progression of numerous types of cancers. To be active, Ras proteins are membrane anchored and organized into nanoclusters, which realize high-fidelity signal transmission across the plasma membrane. Nanoclusters therefore represent potential drug targets. However, targetable protein components of signalling nanoclusters are poorly established.

We previously proposed that the nanocluster scaffold galectin-1 (Gal1) enhances H-Ras nanoclustering by stabilizing stacked dimers of H-Ras and Raf via a direct interaction of dimeric Gal1 with the Ras binding domain (RBD) in particular of B-Raf. Here, we provide further supportive evidence for this model. We establish that the B-Raf preference emerges from divergent regions of the Raf RBDs that were proposed to interact with Gal1. We then identify the L5UR peptide, which disrupts this interaction by binding with low micromolar affinity to the B-Raf-RBD. Its 23-mer core fragment is thus sufficient to interfere with Gal1-enhanced H-Ras nanocluster, reduce MAPK-output and cell viability in *HRAS*-mutant cancer cell lines.

Our data therefore suggest that the interface between Gal1 and the RBD of B-Raf can be targeted to disrupt Gal1-enhanced H-Ras nanoclustering. Collectively, our results support that Raf-proteins are integral components of active Ras nanoclusters.

## Introduction

Ras is a major oncogene and recent advances in its direct targeting have validated its high therapeutic significance ^1, 2^. The three cancer associated Ras genes encode four different protein isoforms, K-Ras4A, K-Ras4B (hereafter K-Ras), N-Ras and H-Ras. These membrane bound small GTPases operate as switchable membrane recruitment sites for downstream interaction partners, called effectors. Downstream of mitogen and growth factor sensing receptors, inactive GDP-bound Ras is activated by guanine nucleotide exchange factors (GEFs), which facilitate GDP/ GTP-exchange ^3, 4^.

The two switch regions of GTP-Ras undergo significant conformational changes upon activation, thus enabling binding to the Ras binding domains (RBDs) of effectors, such as Raf. Current evidence suggests that Ras proteins promiscuously interact with any of the three Raf paralogs, A-, B- and C-Raf. Raf proteins reside as autoinhibited complexes with 14-3-3 proteins in the cytosol and are activated by a series of structural rearrangements that are still not understood in full detail ^5, 6^. The first crucial step is the displacement of the RBD from the cradle formed by the 14-3-3 dimer ^5^. Simultaneous binding of Ras and 14-3-3 to the N-terminal region of Raf is incompatible, due to steric clashes and electrostatic repulsion, which is only relieved if the RBD and adjacent cysteine rich domain of Raf are released from 14-3-3 for binding to membrane-anchored Ras. Allosteric coupling between the N-terminus of Raf and its C-terminus then causes dimerization of the C-terminal kinase domains, which is necessary for their catalytic activity ^6, 7, 8^.

The Ras-induced dimerization of the Raf proteins requires di-/oligomeric assemblies of Ras, called nanoclusters ^9^. Initially it was estimated that 5-20 nm sized nanoclusters contain 6-8 Ras proteins and that nanoclustering was necessary for MAPK-signal transmission ^10, 11, 12^. More recent data revealed that nanoclusters are dominated by Ras dimers ^9, 13^. Intriguingly, Ras nanoclustering can be increased by Raf-ON-state inhibitors that induce Raf dimerization and increase Ras-Raf interaction, suggesting that Raf dimers are integral components of nanocluster ^14, 15^. The reinforced nanoclustering may thus contribute to the paradoxical MAPK-activation that is observed with these inhibitors ^16^.

Currently, less than a dozen proteins are known that can modulate Ras nanoclustering ^17^. These proteins do not share any structural or functional similarities, suggesting that their mechanisms of nanocluster modulation are diverse. The best understood nanocluster scaffold is the small lectin galectin-1 (Gal1), which specifically increases nanoclustering and MAPK-output of active or oncogenic H-Ras ^18, 19, 20^. Consistently, upregulation of galectins has been linked to more severe cancer progression ^21^. For many years, it was mechanistically unclear, how this protein that is best known for binding β-galactoside sugars in the extracellular space affects Ras membrane organization on the inner leaflet of the plasma membrane ^22, 23^. While it was first suggested that the farnesyl tail of Ras is engaged by Gal1 ^24^, it was later on shown that neither Gal1 nor related galectin-3, which is a nanocluster scaffold of K-Ras, bind farnesylated Ras-derived peptides ^25, 26^.

We previously proposed a model of stacked dimers of H-Ras, Raf and Gal1 as the minimal unit of enhanced nanocluster ^27^. We confirmed that Gal1 does not directly interact with the farnesyl tail of Ras proteins, but instead engages indirectly with Ras via direct binding to the RBD of Raf proteins. Given that Gal1 is a dimer, we hypothesized that dimeric Gal1 stabilizes Raf-dimers on active H-Ras nanocluster ^27^. In line with this, in particular B-Raf-dependent membrane translocation of the tumour suppressor SPRED1 by dimer inducing Raf-inhibitors was emulated by expression of Gal1 ^28^. Our mechanistic model suggests that dimeric Gal1 stabilizes the dimeric form of Raf-effectors downstream of H-Ras. This enhances H-Ras/ Raf signalling output, not only by facilitation of Raf-dimerization, but also by an allosteric feedback mechanism that enhances the nanoclustering of H-Ras. Altogether, a transient stacked dimer complex of H-Ras, Raf and Gal1 is formed, which also shifts the H-Ras activity from the PI3K to the MAPK pathway ^27^. However, current galectin inhibitor developments focus on the carbohydrate-binding pocket, which is necessary for its lectin activity in the extracellular space ^29, 30^. Inhibitors that would target its nanocluster enhancement function are missing.

Here we identified a 23-residue peptide that interferes with the binding of Gal1 to the RBD of Raf, thus disrupting H-Ras nanocluster. This peptide validates the Gal1/ RBD interface for future small molecule drug development and supports our model of Gal1-enhanced H-Ras nanoclustering in a stacked-dimer complex.

## Results

### Dimeric Gal1 binds the B-Raf-RBD and stabilizes H-RasG12V nanoclustering

We previously provided evidence that dimeric galectin-1 (Gal1) binds to the Ras binding domain (RBD) of Raf proteins to stabilize active H-Ras nanocluster ^27^ (**Figure 1a**). We first corroborated some features of this stacked-dimer model, using Bioluminesence Resonance Energy Transfer (BRET)-experiments. To this end, interaction partners were tagged with RLuc8 as donor and GFP2 as acceptor and constructs were transiently expressed in HEK293-EBNA (hereafter HEK) cells to monitor the interaction by the increased BRET-signal. In BRET-titration experiments, the characteristic BRET-parameter BRETmax is typically determined. It is a measure for the maximal number of binding sites and the interaction strength, if other interaction parameters, such as complex geometry, are constant ^31^. However, actual binding saturation is typically not reached in cells, and therefore BRETmax cannot be faithfully determined. Hence, we introduced the BRETtop value, which is the maximal BRET-ratio that is reached within a defined range of acceptor/ donor ratios, which is kept constant for BRET-pairs that will be compared ^32^.

**Figure 1.**
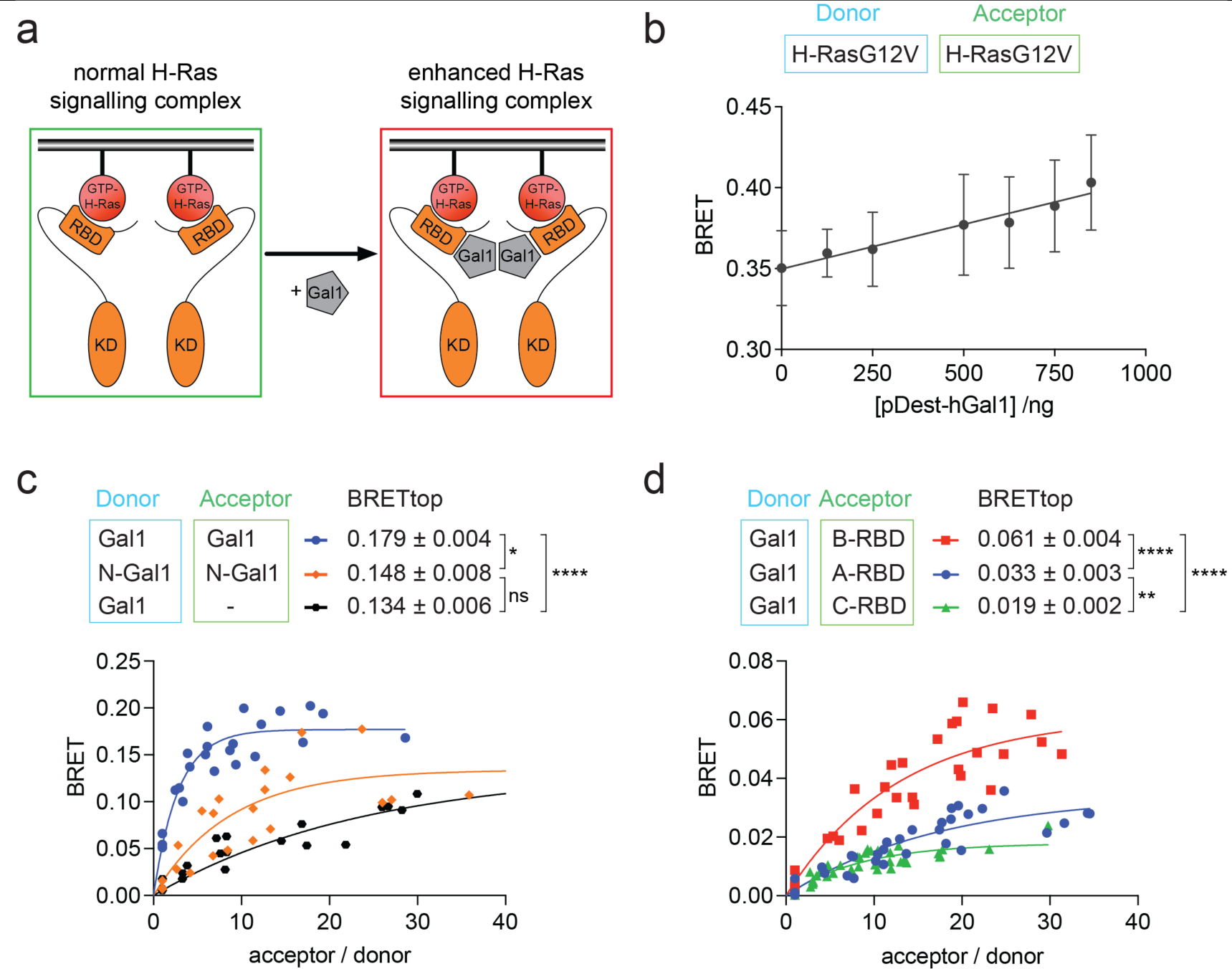
Support for the stacked-dimer model of Gal1-stabilized H-Ras nanocluster. (**a**) Schematic of our model for Gal1 stabilized H-Ras nanocluster. (**b**) Dose-dependent effect of human Gal1 expression (48 h) on H-RasG12V nanoclustering-BRET (donor:acceptor plasmid ratio = 1:5); n = 4. (**c**) BRET-titration curves of the Gal1/ Gal1-interaction as compared to that of dimer-interface mutated N-Gal1. RLuc8-Gal1 was titrated with GFP2 as a control (black); n = 3. (**d**) BRET-titration curves of the Gal1-interaction with the RBDs of A-, B- and C-Raf; n = 3.

In agreement with our earlier results obtained via Förster/ Fluorescence Resonance Energy Transfer (FRET) ^27^, Gal1 expression increased H-RasG12V nanoclustering-BRET in a dose-dependent manner (**Figure 1b**). Mutating four residues at the Gal1 dimer interface (N-Gal1) significantly reduced the BRETtop, confirming that Gal1 is active as a dimer ^33^ (**Figure 1c**). BRET-experiments also confirmed the previously noted interaction preference of Gal1 for B-Raf ^27^ (**Figure S1a**), which was already seen with the RBDs of the corresponding Raf paralogs (**Figure 1d**).

Using computational docking that was based on experimentally determined constraints, we previously proposed a structural model for the binding of Gal1 to the RBD of C-Raf (C-RBD) ^27^ (**Figure S1b**). This model was validated by demonstrating that D113A, D117A mutations in the C-RBD significantly reduced binding to Gal1 ^27^. To further confirm these structural data, we here introduced analogous charge neutralizing mutations D211A and D213A in the B-Raf-derived RBD (B-RBD), and mutation D75A in the A-Raf-derived RBD (A-RBD) (**Figure S1c**). In support of our docking data, the BRETtop of the interaction between Gal1 and either mutant was significantly reduced (**Figure S1d,e**). Consistent with the Raf-paralog specific interaction preference of Gal1, the mutated residues reside in a stretch that is least conserved between the RBDs (**Figure S1c**), which is in agreement with the significant difference in the BRET-interaction data (**Figure 1d**). Taken together with our previously published results ^27^, these data further support our model that Gal1-dimers bind to the RBD in particular of B-Raf, to stabilize the active H-Ras/ Raf stacked-dimer complex and thus an active H-Ras nanocluster.

### Identification of the L5UR-peptide as a disruptor of the Gal1/ RBD interface

Gal1 increases H-Ras-driven MAPK output, and its elevated expression correlates with poorer survival in *HRAS* mutant cancers, such as head and neck squamous cell carcinoma, which frequently displays elevated Gal1 levels ^20, 27^ (**Figure S2a**). Taken together with our H-Ras nanocluster model, these data support targeting of the interface between Gal1 and the RBD as a new strategy against oncogenic H-Ras. We hypothesized that the 52-mer L5UR peptide, which was derived from a Gal1 interaction partner, would be a good starting point for an interface inhibitor. Its residues 22-45 were previously shown to bind with a low affinity (*K*_d_ = 310 µM) to the opposite side of the carbohydrate binding site of Gal1 ^34^. This back-site overlaps with the one we had predicted as RBD-binding site on Gal1 ^27^. We thus expected that the L5UR-peptide would disrupt the Gal1/ RBD interaction and, consequently, the Gal1-augmented H-RasG12V-nanoclustering and MAPK-signalling.

In line with this, expression of untagged L5UR decreased the FRET between mGFP-Gal1 and mRFP-C-RBD in HEK cells (**Figure 2a**). This effect was comparable to the loss observed in the C-RBD-D117A mutant with reduced Gal1-binding (**Figure 2a**) ^27^. For comparison, we tested the effect of Anginex and its topomimetic small molecule analogue OTX-008 ^35^. Anginex is a 33-mer angiostatic peptide that binds to Gal1 at an unknown binding site ^36, 37, 38^. Neither Anginex, nor OTX-008 disrupted the Gal1/ C-RBD interaction as measured by FRET (**Figure 2a**). By contrast, expression of the L5UR-peptide decreased the Gal1-augmented H-RasG12V nanoclustering-FRET. In agreement with previous data ^27^, dimerization-deficient N-Gal1 did not increase nanoclustering-FRET, and co-expression of the L5UR-peptide had no additional effect (**Figure 2b**).

**Figure 2.**
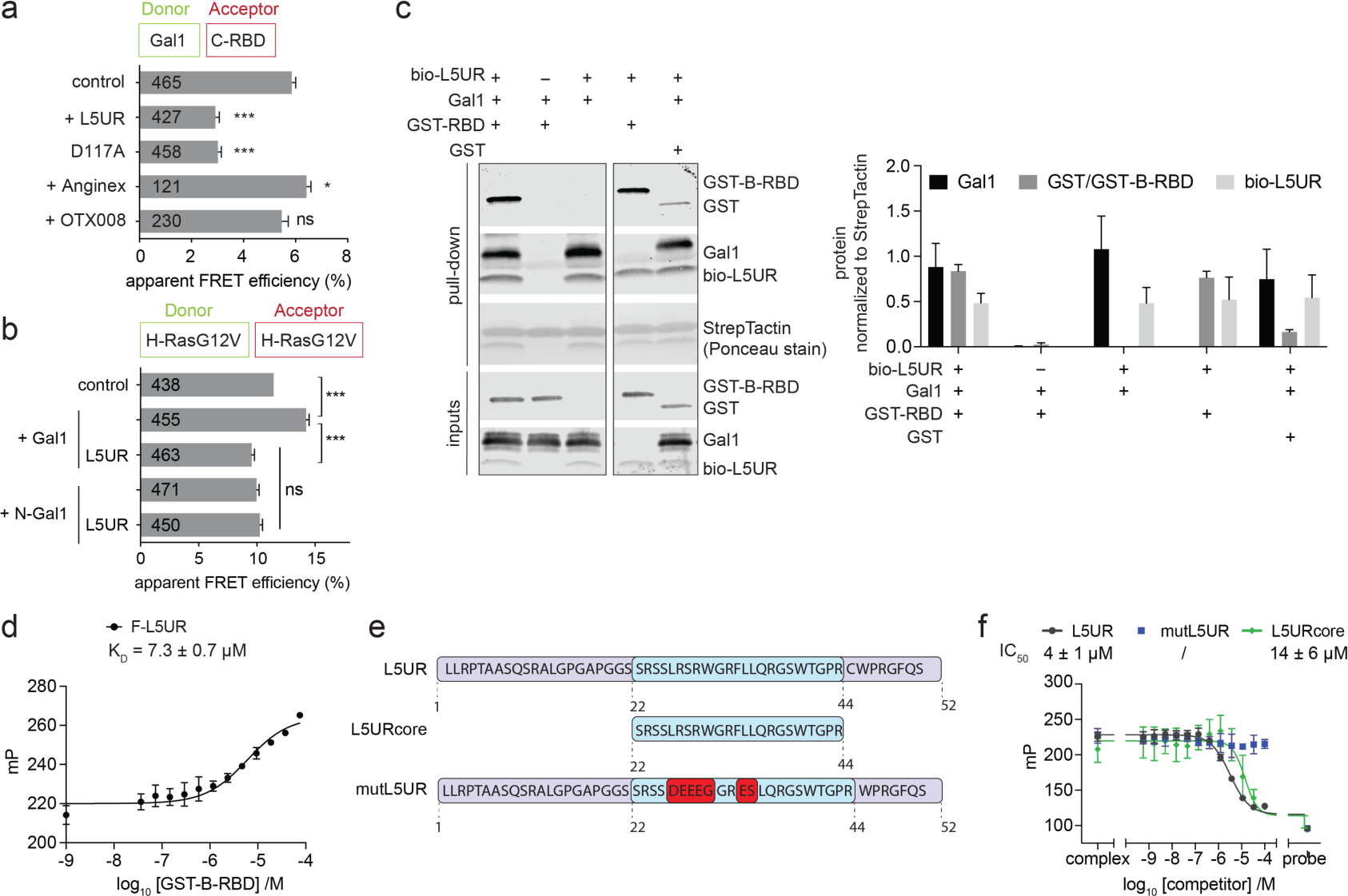
The L5UR-peptide binds to the Raf-RBD and disrupts the Gal1/ RBDcomplex. (**a**) Effect of L5UR expression (24 h) on Gal1/ C-RBD FRET (donor:acceptor plasmid ratio = 1:3); n = 3. (**b**) Effect of L5UR expression (24 h) on Gal1-augmented H-RasG12V nanoclustering-FRET (donor:acceptor plasmid ratio = 1:3); n = 3. (**c**) In vitro pull-down assay with biotinylated L5UR of purified Gal1, GST-B-RBD and GST-only control with example blots (left) and quantification of repeat data (right); n = 3. (**d**) Binding of 10 nM F-L5UR full-length to GST-B-RBD detected in a fluorescence polarization assay; n = 3.

Next, we aimed at confirming that L5UR engages directly with the Gal1/ RBD-interface. We purified His-tagged Gal1 and the GST-tagged B-RBD and performed pulldown experiments with a biotin-tagged L5UR (bio-L5UR) peptide (**Figure 2c**). Interestingly, L5UR pulled down Gal1 and the GST-B-RBD independently from each other (**Figure 2c**). Indeed, fluorescence polarization binding experiments confirmed a micromolar (K_D_ = 7.3 ± 0.7 µM) binding of FITC-tagged full-length L5UR (F-L5UR) to the GST-B-RBD (**Figure 2d**), but not to GST alone (**Figure S2b**). Using a Quenching Resonance Energy Transfer (QRET)-assay, we independently confirmed the micromolar affinity to B-RBD, even with the shortened 22-44 residue core fragment of L5UR labelled with a europium-chelate (Eu-L5URcore) (**Figure S2c**). The L5UR has a high proportion of six positively charged arginine residues in its core region, suggesting that binding of the peptide to the RBD of Raf is predominantly mediated by electrostatic interactions (**Figure S2d**). We therefore introduced seven, mostly charge reversing residue changes in the core-region of the L5UR peptide to generate a non-binding mutant(mutL5UR) (**Figure 2d**). Competitive fluorescence polarization experiments, using F-L5UR as a probe, established that the full-length peptide of L5UR could be displaced from the C-RBD with an IC_50_ = 4 ± 1 µM (**Figure 2f**), and likewise from the B-RBD (**Figure S2e**). As expected, the shorter L5URcore could displace F-L5UR with a slightly reduced potency (IC_50_ = 14 ± 6 µM). Notably, mutL5UR did not reveal any displacement activity in the competitive fluorescence polarization assay (**Figure 2f, Figure S2e**).

In conclusion, L5UR binds with low micromolar affinity to the Raf-RBD. This interaction is lost in the mutL5UR variant, which carries mostly charge-reversal mutations, suggesting that L5UR-binding to the Raf-RBD is driven by electrostatic forces.

### SNAP-tagged L5UR disrupts the Gal1/ B-RBD complex and H-RasG12V nanoclustering in cells

To improve the readout of L5UR-variant expression in cells and eventually enable further functionalization, we designed genetic constructs where a SNAP-tag was added via a long linker to the C-terminus of the peptide (**Figure 3a**). The L5UR-SNAP dose-dependently decreased BRET between Gal1 and the B-RBD to a similar extent as the untagged L5UR, confirming that the SNAP-tag did not increase activity further (**Figure 3b**). In agreement with the binding data (**Figure 2f**), mutL5UR-SNAP did not decrease the BRET signal, nor did the SNAP-tag alone. Immunoblotting confirmed a linear increase of L5UR-SNAP variant expression with increasing amounts of transfected constructs (**Figure 3c,d**).

**Figure 3.**
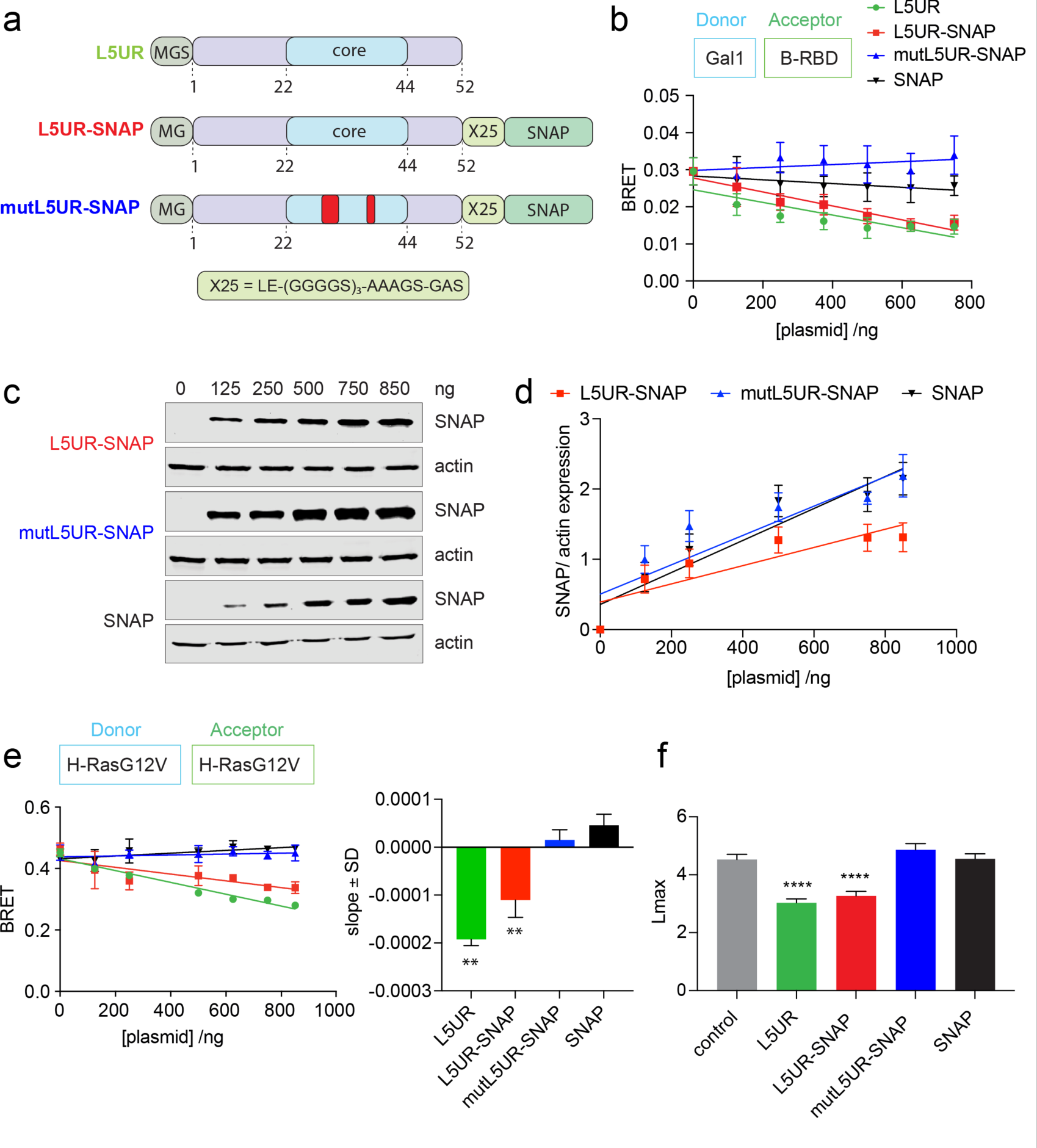
The L5UR and L5UR-SNAP peptides disrupt H-RasG12V nanoclustering. **(a)** Schematics of L5UR derived constructs expressed in cellular assays. The stretch of the core peptide is highlighted in blue, loss-of-function mutations are indicated red. **(b)** Effect of expression of L5UR constructs (48 h) on Gal1/ B-RBD BRET (donor:acceptor plasmid ratio = 1:10); n = 3. (**c,d**) Representative immunoblots (c) and quantification of all repeats (d) showing dose-dependent expression of L5UR constructs (48 h); n = 3. (**e**) Effect of L5UR construct expression (48 h) on H-RasG12V nanoclustering-BRET with co-expression of 200 ng Gal1 (donor:acceptor plasmid ratio = 1:5); n = 2. Statistical comparison was done against the SNAP-only sample. (**f**) Electron microscopy-based analysis of H-RasG12V nanoclustering showing the effects of L5UR-construct expression and controls; n = 15. Higher Lmax values indicate higher nanoclustering.

Consistent with the Gal1/ B-RBD disruption, the L5UR-SNAP construct decreased Gal1-enhanced H-RasG12V nanoclustering-BRET to a similar extent as the untagged L5UR, while again mutL5UR or the SNAP-tag alone had no effect (**Figure 3e**). Neither of these constructs significantly perturbed K-RasG12V nanoclustering-BRET, given that Gal1 is a H-Ras-specific nanocluster scaffold (**Figure S3a**). The disruption of H-RasG12V nanoclustering specifically by L5UR and L5UR-SNAP, but not mutL5UR-SNAP or the SNAP-tag alone, was furthermore confirmed by the classical electron microscopy-based Ras nanoclustering analysis performed on cell membrane sheets (**Figure 3f**). These data therefore confirmed the disruption of H-RasG12V nanoclustering by L5UR- and L5UR-SNAP construct expression.

### TAT-tagged L5UR disrupts MAPK-signalling and inhibits *HRAS*-mutant cancer cell proliferation

Peptides can be rendered cell-permeable by addition of cell penetrating sequences, which facilitate their characterization as prototypic and proof-of-concept reagents ^39^. The 12-residue cell penetrating TAT-peptide that is derived from a Human Immunodeficiency Virus (HIV)-protein, can facilitate cellular peptide uptake ^40, 41, 42^. We therefore chemosynthetically added the TAT-peptide via a PEG-linker to the 23-residue long L5URcore peptide (TAT-L5URcore) and the corresponding loss of function mutant (TAT-mutL5URcore) (**Figure 4a**).

**Figure 4.**
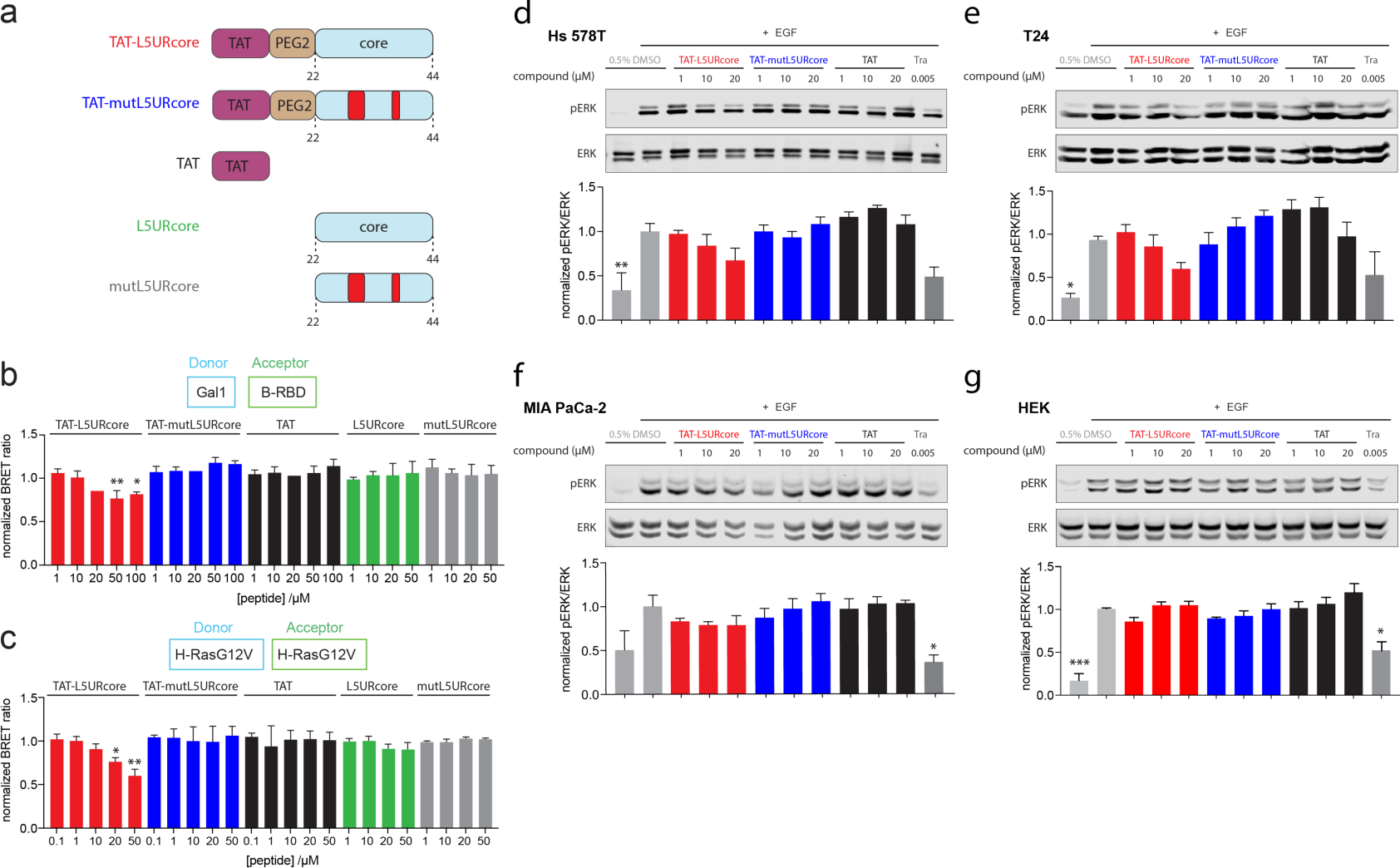
The TAT-tagged L5URcore peptide disrupts Ras-signalling. (**a**) Schematics of L5URcore derived peptides and controls as applied in cellular assays. Loss-of-function mutations of L5UR are indicated in red. Non-TAT peptides are acetylated at the N-terminus. (**b,c**) Effect of cell-penetrating derivatives of L5URcore and control peptides on Gal1/ BRBD BRET (b, donor:acceptor plasmid ratio = 1:10; n = 2) or H-RasG12V nanoclustering-BRET (c, donor:acceptor plasmid ratio = 1:5, co-expression of 200 ng Gal1; n = 3). After 24h expression of plasmids, peptides were added to cells at specified concentrations and incubated for 2 h. (**d-g**) Immunoblot analysis of lysates from Hs 578T (d), T24 (e), MIA PaCa-2 (f) and HEK (g) cells treated with TAT-tagged L5URcore peptides and control compound, trametinib (Tra), for 2 h; n = 4.

To verify cell penetration and on-target activity, we tested the effect of the TAT-peptides in our on-target BRET-assays. Both, the BRET between Gal1 and the B-RBD (**Figure 4b**), as well as H-RasG12V-nanoclustering BRET (**Figure 4c**), were dose dependently disrupted by the TAT-L5URcore peptide. Neither the TAT-peptide alone, nor the mutant TAT-mutL5URcore, or the non-TAT peptides L5URcore and mutL5URcore decreased the BRET-signal in either assay (**Figure 4b,c**).

Based on our model and mechanistic data, signalling and proliferation of *HRAS* mutant cancer cell lines with high Gal1 levels were expected to respond to the nanocluster disrupting TAT-L5URcore peptide. Cancer cell lines Hs 578T (*HRAS-G12D*) and T24 (*HRAS-G12V*), as well as the *KRAS-G12C* mutant MIA PaCa-2 express high levels of Gal1, while HEK cells are devoid of Gal1 (**Figure S3b**). Indeed, treatment of the *HRAS*-mutant cell lines Hs 578T (**Figure 4d**) and T24 (**Figure 4e**) specifically with the TAT-L5URcore peptide reduced cellular pERK levels in a dose-dependent manner, while no such effect was observed in MIA PaCa-2 (**Figure 4f**) or HEK cells (**Figure 4g**).

Consistent with the reduction of MAPK-signalling, the proliferation of the *HRAS*-mutant cancer cell lines Hs 578T (**Figure 5a,e**) and T24 (**Figure 5b,e**) was significantly reduced by TAT-L5URcore, but not the control TAT-peptides. However, this time also proliferation of MIA PaCa-2 (**Figure 5c,e**) and HEK cells (**Figure 5d,e**) was affected as revealed by our normalized area under the curve DSS3-analysis, where a higher DSS3-score corresponds to a higher anti-proliferative activity (Figure 5e).

**Figure 5.**
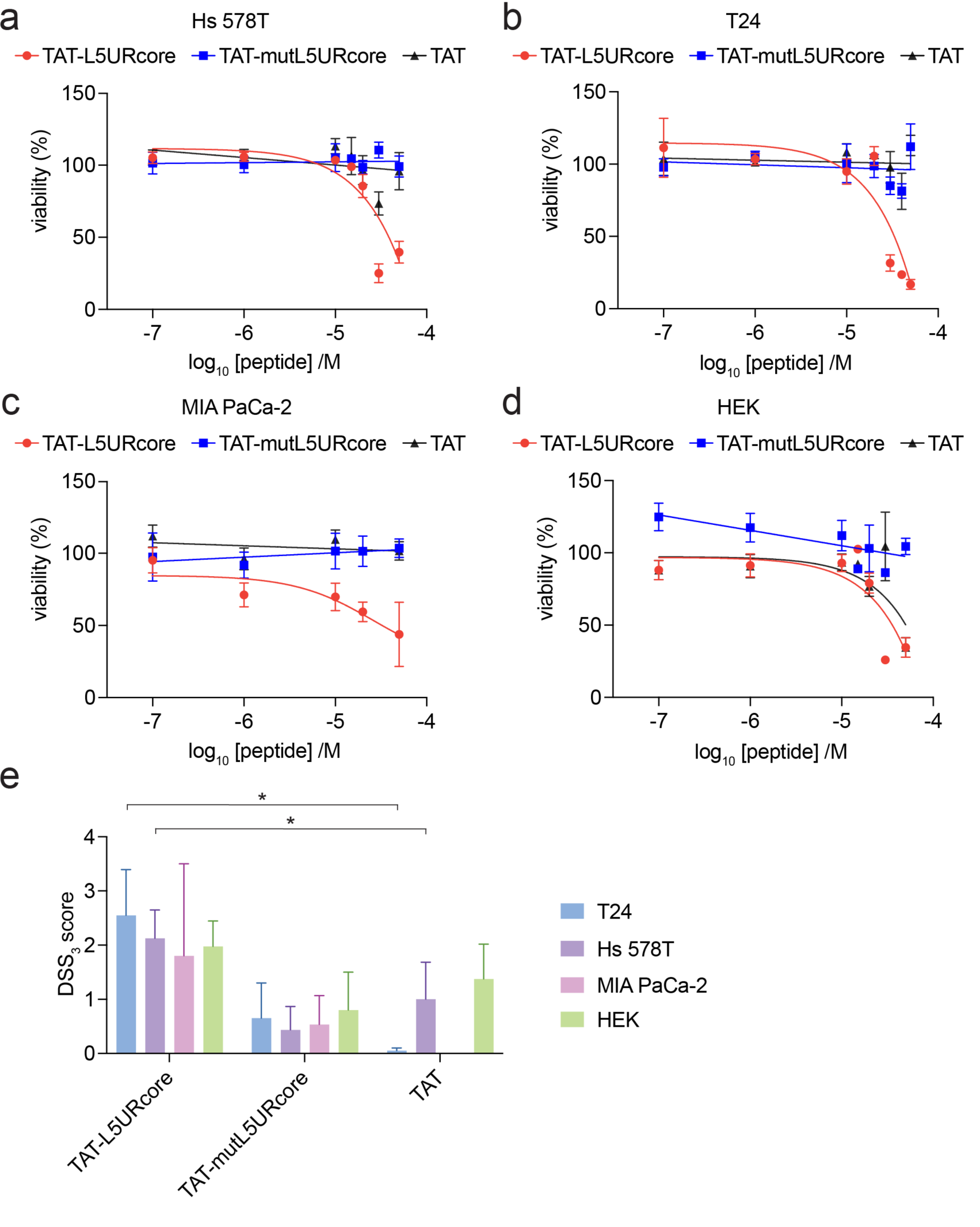
*HRAS*-mutant cancer cell proliferation is decreased by TAT-L5UR peptides. (**a-d**) 2D cell viability of Hs 578T (a), T24 (b), MIA PaCa-2 (c) and HEK (d) cells in response to 48 h treatment with TAT-L5URcore peptides and TAT-control; n = 3. (**e**) Drug sensitivity score (DSS_3_), an area under the curve metric, calculated for the viability data in (a-d). A higher value indicates a stronger anti-proliferative effect. TAT-control was used as a reference for statistical comparisons.

This broader effect on cell proliferation may indicate that the TAT-L5URcore interferes also with other signalling pathways than the MAPK-pathway that are relevant for cell proliferation.

## Discussion

We here demonstrate that the 23-residue L5URcore peptide binds with micromolar affinity to the Raf-RBD thus disrupting the interaction with Gal1. The peptide can therefore interfere with Gal1-enhanced nanocluster of active H-Ras, MAPK-signalling and cell proliferation of *HRAS* mutant cancer cell lines. The activity of this peptide validates the importance of the Gal1/ Raf interaction in Gal1-stabilized H-Ras nanocluster and indirectly supports our stacked dimer model.

However, several questions remain unanswered. For instance, it is currently unknown how Gal1 positively regulates H-Ras nanocluster, but negatively K-Ras nanocluster ^27^. Vice versa, how the related galectin-3 (Gal3) increases specifically K-Ras nanocluster is not known ^43, 44, 45^. In the context of our stacked-dimer model, it is conceivable, that galectins stabilize specific Raf-dimers and thus nanoclustering of specific Ras isoforms. Indeed, Gal1 distinguishes between the RBDs from A-, B-, and C-Raf and most strongly engages the B-Raf-RBD. For K-Ras evidence exists that it binds preferentially with B-/ C-Raf-dimers ^14, 46^, while for Gal1 augmented H-Ras nanocluster our previous data suggested a particular relevance for B-/ A-Raf dimers ^27^. One would therefore predict that these dimers are specifically stabilized by Gal3 and Gal1, respectively. However, it is not entirely plausible how symmetrical dimers of galectins, or in the case of Gal3 potentially even oligomers ^23^, would stabilize asymmetric dimers of Raf proteins. Heterodimerization of galectins could provide a solution to this problem. In humans, 15 different galectins are found and only Gal1 and Gal3 are characterized as nanocluster scaffolds so far ^23^. Given the relatedness in this protein family, it is plausible to assume that other galectins have a similar activity and potentially mixed galectin-dimers could form that then stabilize the asymmetric dimers of Raf. Therefore, a complex equilibrium of mixed oligomers that partly stabilize and partly compete and sequester could be the answer to the intricate problem of Ras-isoform specific nanoclustering effect of galectins.

The TAT-L5URcore provides a unique tool to investigate the functioning of Ras nanocluster further. In contrast to current galectin inhibitors, which target the carbohydrate-binding pocket ^29, 30^, the L5UR-peptide acts via a novel mode-of-action that exploits galectin’s nanocluster stabilizing activity. The intermediate size below 3 kDa of the TAT-L5URcore peptide represents a relevant starting point for the development of smaller molecules with analogous mode-of-action. The properties of this peptide and the putative target site suggest that not a distinct pocket, but an assembly of charge interactions are currently the major driving force for its affinity. Regarding size and mechanism of action, L5URcore contrasts to the NS1-monobody, which specifically binds to the allosteric lobe of K-Ras and H-Ras to disrupt nanoclustering ^47^. Given the size of the monobody of ∼10 kDa it is likely that the steric hindrance caused by this large ligand is mostly responsible for the interference with nanoclustering. With the identification of the targetable site on the Raf-RBD and with more insight into the structure of the Gal1/ RBD complex, it will be possible to identify improved binders with higher affinity and specificity in the future. Both competitive screening as well as structure-based design of peptidomimetics present opportunities for future improvements.

Targeting of the augmenting effect of Gal1 on H-Ras nanoclustering is quite different from approaches focusing on the main nodes of the Ras-MAPK-pathway. Both mechanistic and genetic evidence suggest that Gal1 acts as a positive modifier that is associated with a worse progression of *HRAS* mutant cancers, notably head and neck cancers that are frequently associated with high Gal1 levels (**Figure S2a**). While *HRAS* is overall the least frequently mutated *RAS* gene (in 1.3 % of cancer patients), it is mutated in > 5% of head and neck squamous cell carcinomas (HNSCC) ^48^. Prognosis for patients with recurrent and metastatic HNSCC is still poor ^49^. While tipifarnib, a farnesyltransferase inhibitor shows promising efficacy in HNSCC patients, there is still a need for potent treatments ^50^. By interfering with the interface of Gal1 and Raf-proteins, one does not eliminate other functions of these proteins and therefore may specifically achieve a normalization of the signalling activity. This would be beneficial in regard to side effects, as normal tissue functions could continue to progress. We expect that our L5UR peptide work will provide new perspectives on how to target Ras nanocluster.

## Materials and methods

### Expression Constructs

Here we refer to the 52-mer fragment derived from residues 38-89 of the unique region of the λ5-chain (λ5-UR) of the pre-B-cell receptor as L5UR. This unique region bears no similarity to known proteins ^34^. The pClontech-L5UR was made by excising L5UR cDNA from pET28a-L5UR (gift from Dr. Elantak), using NheI – XhoI sites and subcloned into pmCherrry-C1 (Clontech, #632524). This removed the mCherry cDNA from the expression vector leaving only the full-length L5UR. Vector pcDNA-Hygro-Anginex was a gift from Prof. Thijssen ^38, 51^. Expression clones were mostly produced by multi-site gateway cloning as described in our previous studies ^32, 52, 53^. Some expression clone genes were synthesized and cloned into desired vectors by the company GeneCust, France. A list of all the clones used in the study and their sources are given in **Table S1**.

### Cell Culture

Hs 578T, T24, MIA PaCa-2 and BHK-21 cells were obtained from DSMZ-German Collection of Microorganisms and Cell Cultures GmbH or ATCC. HEK293-EBNA cells were a gift from Prof. Florian M. Wurm, EPFL, Lausanne. All cell lines were cultured in a humidified incubator maintained at 37 °C and 5 % CO_2_, in Dulbecco’s modified Eagle Medium (DMEM) (Gibco, #41965039) supplemented with 9 % v/v Fetal Bovine Serum (FBS) (Gibco, #10270106), 2 mM L-Glutamine (Gibco, #25030081) and penicillin-streptomycin (Gibco, #15140122) 10,000 units/ mL (complete growth medium), in T75 culture flasks (Greiner, #658175). Cells were regularly passaged 2-3 times a week and routinely tested for mycoplasma contamination using MycoAlert Plus mycoplasma Detection kit (Lonza, #LT07-710).

### Bacterial strains

Competent E. coli BL21 Star (DE3)pLysS and E. coli DH10B were grown in Luria-Bertani (LB) medium (Sigma, #L3022) at 37 °C, with appropriate antibiotics unless otherwise stated.

### Protein purification

For protein expression, a 16 h culture was set by inoculating colonies into appropriate volume of antibiotic-supplemented LB media incubated 16 h at 37 °C. The next day, 25 mL of the culture was added to 1 L of LB and incubated at 37 °C until OD at 600 nm reached 0.6-0.9, at which point protein expression was induced by adding isopropyl β-D-1-thiogalactopyranoside (IPTG) (VWR, #437145X) at the final concentration of 0.5 mM. GST-tagged B-Raf-RBD (residues 155-227 of B-Raf) and GST-tagged C-Raf-RBD (residues 50-134 of C-Raf) protein expression was induced for 4 h at 23 °C and the His-tagged protein expression was induced for 16 h at 25°C. Afterwards the cell pellet was collected by centrifugation, rinsed in PBS and stored at −20 °C until purification.

For GST-tagged protein purification, cells were lysed by resuspending the pellet in a buffer consisting of 50 mM Tris-HCl pH 7.5, 150 mM NaCl, 2 mM DTT, 0.5 % v/v Triton-X 100, 1× Protease Inhibitor Cocktail (Thermo Scientific Pierce Protease Inhibitor Mini Tablets, EDTA-free, #A32955) and by sonication on ice using a Bioblock Scientific Ultrasonic Processor instrument (Elmasonic S 40 H, Elma). Lysates were cleared by centrifugation at ∼18,500 ×g for 30 min at 4 °C. For GST-tagged proteins, the cleared lysate was incubated with 500 µL glutathione agarose slurry (GE Healthcare, #17-0756-01) (resuspended 1:1 in lysis buffer) for 3 h at 4 °C with gentle rotation. Next, the supernatant was removed, and beads were washed five times with 1 mL of washing buffer consisting of 50 mM Tris-HCl at pH 7.5, 500 mM NaCl, 5 mM DTT, 0.5 % (v/v) Triton-X 100. Next, beads were rinsed three times with 1 mL of equilibration buffer (50 mM Tris-HCl pH 7.5, 150 mM NaCl, 2 mM DTT). GST-tagged protein was eluted off the beads by using 20 mM glutathione solution (Sigma-Aldrich, #G4251-5G). Fractions were analyzed by resolving on 4-20 % gradient SDS-PAGE (BioRAD #4561094 or #4651093), stained with Roti Blue (Carl Roth Roti-Blue quick, #4829-2) and dialyzed into a final dialysis buffer (50 mM Tris-HCl at pH 7.5, 150 mM NaCl, 2 mM DTT, 10 % (v/v) glycerol) by using a D-Tube Dialyzer with MWCO 6-8 kDa (Millipore, #71507-M) for 16 h at 4 °C. Protein concentration was measured using NanoDrop 2000c Spectrophotometer (Thermo Fischer Scientific) and stored at −80 °C.

For GST-tag removal, the cleared lysate was incubated with 500 µL of glutathione agarose slurry (resuspended 1:1 in lysis buffer) for 5 h at 4 °C with gentle rotation, then proceeded to washing steps as described above. The beads were rinsed with equilibration buffer and then with dialysis buffer before the excess was drained as much as possible. The beads were then resuspended in 650 µL of dialysis buffer and 100 U of Thrombin (GE Healthcare, #GE27-0846-01), to a final volume of 1 mL. The next day, the untagged protein was collected by applying supernatant to 1 mL polypropylene column and the flow-through was collected as fraction 1. The beads were washed once more with 1 mL of dialysis buffer and the flow-through was collected as fraction 2. The two fractions were analysed by resolving on 4-20 % gradient SDS-PAGE and stained with Roti Blue. Protein concentration was measured using NanoDrop and stored at −80 °C.

For His-tagged protein purification, the cells were resuspended in lysis buffer (50 mM Tris-HCl at pH 7.4, 150 mM NaCl, 5 mM MgSO_4_, 4 mM DTT, 100 mM β-lactose, 100 μM phenylmethylsulfonyl fluoride) with ∼ 5 mg of DNAseI (Merck, #10104159001) and ∼ 5 mg of lysozyme (Thermo Fisher Scientific, #89833). Cells were lysed using a LM10 microfluidizer (Microfluidics, USA) at 18000 PSI and cell debris were separated by centrifugation (4 °C, 30 min, 75,600 ×g, JA25.50 rotor Beckman Coulter). The supernatant was loaded on an affinity chromatography column (GE Healthcare, His-Trap FF crude, #17-5286-01) with a flow rate of 1 mL/ min. A total amount of 10 CV (column volumes) 10 % elution buffer (50 mM Tris-HCl pH 7.4, 150 mM NaCl, 5 mM MgSO_4_, 100 mM β-lactose, 4 mM DTT, 1 M Imidazole) and 90 % lysis buffer (50 mM Tris-HCl pH 7.4, 150 mM NaCl, 5 mM MgSO_4_, 4 mM DTT, 100 mM β-lactose) with a flow rate of 2 mL/ min was applied. The protein was then eluted using 5 CV of elution buffer (50 mM Tris-HCl pH 7.4, 150 mM NaCl, 5 mM MgSO_4_, 100 mM β-lactose, 4 mM DTT, 1 M Imidazole). Afterwards, the protein was injected into a size exclusion chromatography system (GE Healthcare, HiLoad 16/600 Superdex 75 pg, #28-9893-33) using SEC buffer (20 mM HEPES pH 7.4, 150 mM NaCl, 5 mM MgSO_4_, 100 mM β-lactose, 4 mM DTT) and a flow rate of 1 mL/ min. Protein containing fractions were pooled, concentrated (MWCO = 3 kDa) to 16.1 mg/ mL, snap frozen in liquid nitrogen and stored at −80 °C. The protein concentration was measured using NanoDrop 2000c Spectrophotometer (Thermo Fisher Scientific).

### Fluorescence polarisation assays

The fluorescence polarisation assay was adapted from our previously established protocol ^52, 54^. The non-labelled L5UR and their derivatives and FITC-labelled peptides were obtained from Pepmic Co., China. F-L5UR was synthesised by attaching fluorescein to the N-terminus amino group, leucine of L5UR peptide via aminohexanoic acid linker.

For the direct binding assay, the GST-B-RBD or GST, was 2-fold diluted in an assay buffer composed of 50 mM Tris HCl pH 7.4, 50 mM NaCl, 5 mM DTT and 0.005 % v/v Tween 20 in a black low volume, round bottom 384-well plate (Corning, #4514). Then 10 nM F-L5UR peptide was added to each well and incubated for 20 min at ∼22 °C on a horizontal shaker. The fluorescence polarisation measurement was performed on the Clariostar (BMG Labtech) plate reader, using a fluorescence polarization module (λ_excitation_ 482 ± 8 nm and λ_emission_ 530 ± 20 nm). The fluorescence intensity signal was recorded from vertical (I_v_)- and horizontal (I_h_)-polarized light. The milli fluorescence polarisation, mP, was determined from the measured fluorescence intensities, calculated according to,

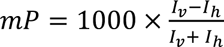

where I_v_ and I_h_ are the fluorescence emission intensities detected with vertical and horizontal polarization, respectively. The mP was plotted against concentration of the GST-RBD and the K_D_ value of the F-L5UR was calculated using a quadratic equation,

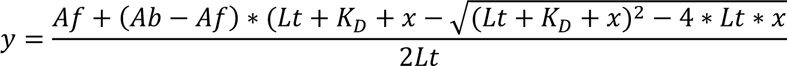

Af is the anisotropy value of the free fluorescent probe, Ab is the anisotropy value of the fluorescent probe/ protein complex, Lt is the total concentration of the fluorescent probe, K_D_ is the equilibrium dissociation constant, x is total concentration of protein and y is measured anisotropy value ^32, 54^. K_D_ is measured in the same unit as x.

For competitive fluorescence polarisation experiments, the non-labelled peptides were three-fold diluted in the assay buffer and then a complex of 5 nM F-L5UR peptide and 200 nM B-RBD was added to the dilution series to a final volume of 20 µL per well in 384-well plate. After 30 min incubation at ∼22 °C, the fluorescence polarisation was read. The logarithmic concentration of peptide was plotted against the mP value and the data were fit into log (inhibitor) vs response four parameters equation in GraphPad, and the IC50 values were derived. IC_50_ values were converted into K_D_ values as described earlier ^55^.

### QRET assays

The QRET assays were modified from our previously described quenching luminescence assays ^56, 57, 58^. Ac-K-L5URcore was conjugated with nonadentate europium chelate, {2,2’,2",2’"-{[4’-(4’"-isothiocyanatophenyl)-2,2’,6’,2"-terpyridine-6,6"-diyl]bis(methylene-nitrilo)}tetrakis(acetate)}europium(III) (QRET Technologies, Finland) via the epsilon amine of the N-terminal lysine that was added to the L5UR-core peptide sequence and purified with analytical reverse-phase HPLC.

The current homogeneous QRET binding assay is based on the quenching of non-bound Eu-K-L5URcore with MT2 quencher (QRET Technologies), while bound labelled peptide is luminescent. In the assay, B-RBD was 2-fold diluted in an assay buffer containing 10 mM HEPES pH 7.4, 10 mM NaCl added in 5 μL to a white low volume, round bottom 384-well plate. Eu-K-L5UR core peptide (29 nM), mixed with MT2 according to the manufacturer’s instructions in the assay buffer supplemented with 0.01 % (v/v) Triton X-100, was added in 5 μL volume to wells, and incubated for 30 min at ∼22 °C on a shaker. The luminescence was measured with Tecan Spark multimode microplate reader (Tecan, Austria) in time-resolved mode using λ_excitation_ 340 ± 40 nm and λ_emission_ 620 ± 10 nm with 800 μs delay and 400 μs window times.

### In vitro pull-down assays with recombinant proteins

Biotinylated L5UR (Bio-L5UR) peptide was synthesised as described above. GST-B-Raf-RBD (155-227), His-Gal1, His-N-Gal1 and GST were prepared as described above. Each protein in the assay was used at 2 µM concentration and the peptide was at 4 µM. Volume of the reaction was 150 µL. First, peptide and Gal1 were pre-incubated for 30 min at 37 °C, then GST-B-RBD or GST alone were added, and the reaction continued for another hour. Control reaction mixes contained DMSO instead of the peptide. At the end of the reaction time, 10 µL of each sample was withdrawn for SDS-PAGE analysis as inputs. For pull-downs, 5 µL of the beads were taken per sample. To prepare the beads, appropriate volume of the slurry was pipetted into 15 mL falcon tubes and centrifuged at 830 ×g for 1 min to remove ethanol-containing supernatant. The falcon tube was topped up to 15 mL with distilled water and centrifugated for 1 min to remove water. This washing step was repeated three times. Finally, the beads were resuspended in distilled water so that the final bead volume was 4 × diluted and, therefore, 20 µL were pipetted to each tube. Pull-down was conducted by incubating samples on a rotating wheel at room temperature (20-25 °C) for 1 h. Then, the samples were centrifuged for 1 min at 830 ×g at 4 °C. The supernatant was discarded, the beads were rinsed with 250 µL of washing buffer (50 mM Tris HCl pH 7.5, 150 mM NaCl, 4 mM β-mercaptoethanol, 0.05 % (v/v) NP-40, 10 % (v/v) Glycerol) for the total of 1 h at 4 °C, with four exchanges of the washing buffer. The bound material was eluted off the beads by adding 2 × SDS-PAGE sample buffer and incubating for 5 min at 95 °C. The analysis was done by resolving the samples (8 µL of the input samples and 10 µL of the eluted material) on 4-20 % gradient SDS-PAGE gels and analysed by Western blotting. A list of all the antibodies used in the study and their sources are given in **Table S1**.

### Electron microscopic analysis of Ras-nanoclustering

To quantify the nanoclustering of a component integral to the plasma membrane (PM), the apical PM sheets of baby hamster kidney (BHK) cells expressing a GFP-tagged H-Ras construct were fixed with 4 % (w/v) PFA and 0.1 % (w/v) glutaraldehyde. GFP anchored to the PM sheets was probed with 4.5 nm gold particles pre-coupled to anti-GFP antibody. Following embedment with methyl cellulose, the PM sheets were imaged using transmission electron microscopy (JEOL JEM-1400). Using the coordinates of every gold particle, the Ripley’s K-function calculated the extent of nanoclustering of gold particles within a selected 1 μm^2^ PM area:

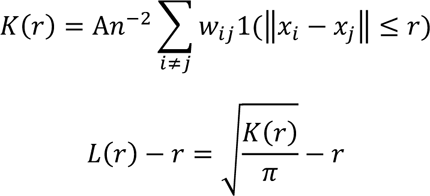

where *n* gold particles populate in an intact area of *A; r* is the length between 1 and 240 nm; || · || indicates Euclidean distance where 1(·) = 1 if ||*x_i_*-*x_j_*|| ≤ r and 1(·) = 0 if ||*x_i_*-*x_j_*|| > r; *K(r*) specifies the univariate K-function. *w_ij_*^-1^ is a parameter used for an unbiased edge correction and characterizes the proportion of the circumference of a circle that has the center at *x_i_* and radius ||*x_i_*-*x_j_*||. Monte Carlo simulations estimates the 99 % confidence interval (99 % C.I.), which is then used to linearly transform *K*(*r*) into *L*(*r*) – *r*. On a nanoclustering curve of *L*(*r*) – *r* vs. *r,* the peak *L*(*r*) – *r* value is used as summary statistics for nanoclustering and is termed as *Lmax.* For each condition, at least 15 PM sheets were collected for analysis. To analyse statistical significance between conditions, bootstrap tests compare our point patterns against 1000 bootstrap samples.

### Immunoblotting

Routinely, 4–20 % Mini-PROTEAN TGX Precast Protein Gels, 10-well, 50 µL, or 30 µL (BioRad, #4561094 or #4651093) were used, unless stated otherwise. For protein size reference, All Blue (Precision Plus Protein All Blue Prestained Protein Standards (BioRad, #1610373) or Page Ruler Prestained (Thermo Fisher Scientific, #26616) were used.

For ERK activity studies, Hs 578T, T24, MIA PaCa-2 and HEK cells were grown in a 6-well plate for 24 h. After 16h serum starvation, the cells were treated for 2 h with the L5UR derived TAT-peptides or DMSO control, before they were stimulated with 200 ng/ mL EGF for 10 min. The cell lysates were then prepared using a buffer composed of 150 mM NaCl, 50 mM Tris-HCl pH 7.4, 0.1 % (w/v) SDS, 1 % (v/v) Triton X-100, 1 % (v/v) NP40, 1 % (w/v) Na-deoxycholate, 5 mM EDTA pH 8 and 10 mM NaF completed with 1 × protease inhibitor cocktail (Pierce, #A32955) and 1 × phosphatase inhibitor cocktail (Roche PhosSTOP, #490684001). The total protein concentration was determined using Bradford assay (Protein Assay Reagent, BioRad, #5000006) and 25 µg cell lysate was loaded on a 10 % homemade SDS-PAGE gel.

For immunoblotting, gels were transferred onto 0.2 µm pore–size nitrocellulose membrane by using Trans-Blot Turbo RTA Midi 0.2 µm Nitrocellulose Transfer Kit, for 40 blots (BioRad, #1704271). The membranes were blocked with TBS or PBS with 0.2 % (v/v) Tween20 and 2 % BSA. Primary antibodies were incubated at 4 °C for 16 h or for 1-3 h at room temperature (20-25 °C). All secondary antibodies were diluted at 1:10,000 in a blocking buffer and were incubated for 1 h at room temperature (20-25 °C). A detailed list of all the antibodies used in the study and their sources are given in **Table S1**.

### Fluorescence Lifetime Imaging Microscopy (FLIM)-FRET analysis

FLIM-FRET experiments were conducted as described previously ^27, 59, 60^. About 120,000 HEK cells were seeded per well in a 6-well plate (Greiner, #657160) with a cover slip (Carl Roth, #LH22.1) and grown for 18 to 24 h. For H-RasG12V nanoclustering-FRET, the cells were transfected with a total of 1 µg of mGFP/ mCherry-tagged H-RasG12V at a donor (D):acceptor (A)-plasmid ratio of 1:3. In addition, 0.75 µg of other plasmids encoding L5UR, Gal1 or N-Gal1 were co-transfected. For Gal1/ C-RBD FRET-interaction, the cells were transfected with 2 µg mGFP-rtGal1 and mRFP-C-RBD (D:A, 1:3) or mGFP-rtGal1 and mRFP-C-RBD-D117A pair (D:A, 1:3). In addition, cells were co-transfected with 1.5 µg pClontech-C-L5UR, the pcDNA-Hygro-Anginex or compound OTX008 (Cayman Chemicals, #23130). All transfections were done using jetPRIME (Polyplus, #114-75) transfection reagent according to the manufacturer’s instructions. After 4 h of transfection the medium was changed. The next day, the cells were fixed with 4 % w/v PFA. The cells were mounted with Mowiol 4-88 (Sigma-Aldrich, #81381). An inverted microscope (Zeiss AXIO Observer D1) with a fluorescence lifetime imaging attachment (Lambert Instruments) was used to measure fluorescence lifetimes of mGFP. fluorescein (0.01 mM, pH 9) was used as a fluorescence lifetime reference (τ = 4.1 ns). Averaged fluorescence lifetimes were used to calculate the apparent FRET efficiency as described ^59, 60^.

### BRET assays

We employed the BRET2 system where RLuc8 and GFP2 luminophores were used as the donor and acceptor, respectively, with coelenterazine 400a as the substrate. A CLARIOstar plate reader from BMG Labtech was used for BRET and fluorescence intensity measurement. The BRET protocol was adapted as described by us ^61^.

In brief, 150,000 to 200,000 HEK293-EBNA cells were seeded per well of a 12-well plate (Greiner Bio-One, #665180) and grown for 24 h in 1 ml of complete DMEM. The next day, the cells were transfected with ∼ 1 µg of plasmid DNA per well using 3 µL jetPRIME transfection reagent. For the donor saturation titration, 25 ng of the donor plasmid was transfected with an acceptor plasmid concentration ranging from 25 ng to 1000 ng. pcDNA3.1(-) (Thermo Fisher Scientific, #V79520) was used to normalize the amount of DNA per well. 48 h after transfection, cells were collected in PBS and plated in a white 96-well plate (Nunc, Thermo Fisher Scientific, #236108).

First the fluorescence intensity of GFP2 was measured (λ_excitation_ 405 ± 10 nm and λ_emission_ 515 ± 10 nm), which is directly proportional to the acceptor expression (RFU). Then 10 µM of coelenterazine 400a (GoldBio, #C-320) was added to the cells and BRET readings were recorded simultaneously at λ_emission_ 410 ± 40 nm (RLU) and 515 ± 15 nm (BRET signal). Emission intensity measured at 410 nm is directly proportional to the donor expression. Raw BRET ratio was calculated as the ratio of BRET signal/ RLU. Background BRET ratio was obtained from cells expressing only the donor. Background BRET ratio was subtracted from raw BRET ratio to obtain the BRET ratio, plotted labelled as BRET. The expression was calculated as the ratio of RFU/RLU. The relative expression, acceptor/ donor, plotted in the x-axis in corresponding figures, was obtained by normalizing RFU/RLU values from cells transfected with equal dose of donor and acceptor plasmids ^46^.

The BRET ratio and acceptor / donor from various biological repeats were plotted together and the data were fit into a hyperbolic equation in Prism (GraphPad). The one phase association equation of Prism 9 (GraphPad) was used to predict the top asymptote Ymax-value, which was taken as the BRETtop. The BRETtop value represents the top asymptote of the BRET ratio reached within the defined acceptor / donor range.

For the dose-response BRET assays, the donor and acceptor plasmid concentration were kept constant, as indicated in the corresponding figure legends. HEK293-EBNA cells were grown in 12-well plate for 24 h in complete DMEM and next day, donor and acceptor plasmids were transfected along with modulator plasmid ranging from 125 ng to 850 ng. After 48 h of expression the cells were collected in PBS and BRET measurements were carried out.

For treatment with peptides, HEK cells were batch transfected. After 24 h of transfection, cells were re-plated in white 96-well plate in phenol red-free DMEM. After another 48 h, peptides were added to cells at concentration ranging from 0.1 µM to 100 µM. After 2 h incubation at 37 °C, the plate was brought to room temperature (20-25 °C) before taking BRET measurements as indicated above. The concentration of the transfected L5UR-modulator plasmid or applied peptide was plotted against the BRET-value and the data were fit into a straight-line equation using Prism.

### Cell Viability Assay and Drug Sensitivity Score (DSS) Analysis

The cells were seeded in low attachment, suspension cell culture 96-well plates (Greiner, #655185). About 2000 T24, MIA PaCa-2 and HEK cells and 5000 Hs 578T cells were seeded per well in 50 µL complete growth medium. 24 h later the cells were treated with 50 µL 2 × peptide diluted in growth medium, or 0.2 % (v/v) of the positive control, benzethonium chloride (Sigma-Aldrich, #B8879). 48 h after the peptide treatment 10 % (v/v) of alamarBlue reagent (Thermo Fisher Scientific, #DAL1100) was added to each well and incubated for 4 h at 37°C. Using a CLARIOstar plate reader the fluorescence signal (λ_excitation_ 560 ± 5 nm and λ_emission_ 590 ± 5 nm) was recorded. The florescence signal was normalized against the negative control, here DMSO, representing 100 % viability. Additionally, the data was analysed using Breeze 2.0 to determine a drug sensitivity score (DSS), a normalized area under the curve (AUC). Here we plot only one of the output values from the Breeze pipeline ^62^, the DSS_3_ value, which was calculated as

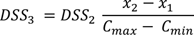

where DSS_2_ is given by the equation 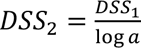

and DSS_1_ is given by the equation 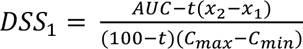

### Statistical analysis

Data were analyzed using Graph Pad prism 9.0 software. The number of independent biological repeats (n) for each dataset is provided in the figure legends.

If not stated otherwise means and standard errors (SEM) are plotted. All BRETtop data were compared using the extra sum-of-squares F test. All other statistical analyses were performed using One-way ANOVA. A p-value of < 0.05 was considered statistically significant and the statistical significance levels were annotated as: * = P < 0.05; ** = P < 0.01; *** = P < 0.001; **** = P < 0.0001, or ns = not significant.

## Data availability

All relevant data supporting this study are available within the manuscript and supplementary data. Source data are provided with the manuscript. All unique/ stable reagents generated in this study are available from the corresponding author with a completed materials transfer agreement. This study did not report standardized datatypes.

## Acknowledgements

The study was supported by the grant INTER/NWO/19/14061736-HRAS-PPi of the Luxembourg National Research Fund (FNR) and the Dutch Research Council (NWO) to DA and TG, as well as FNR-grant INTER/Mobility/2021/BM/15591725/panRAFi-PB to GM. Prof. Marc Therrien (IRIC, Université de Montréal, Canada) is gratefully acknowledged for hosting GM in his laboratory. Dr. Hugo Lavoie and Dr. Ting Jin (IRIC, Université de Montréal, Canada) are thanked for their support and advice to GM during his sabbatical. Geneviève Arseneault (IRIC, Université de Montréal, Canada) is thanked for the technical support. pET28a-L5UR was a gift from Dr. Latifa Elantak (CNRS Marseille, France). pcDNA-Hygro-Anginex was a gift from Prof. Victor L. Thijssen (VU Amsterdam, Netherlands).

## Author Contributions

DA and TG conceived the study.

GM collected and evaluated BRET and FP data.

CS collected and evaluated BRET, FP, signaling and cell viability data and purified proteins.

AYV synthesized all of the peptides.

KP purified proteins and performed and evaluated pull-down experiments.

MK collected and evaluated FLIM-FRET data.

YZ and NA collected and evaluated EM-nanoclustering data.

HH collected and evaluated QRET data.

AG performed bioinformatics analysis of survival and cancer type frequency.

TG and DA jointly supervised the study.

GM, CS, AYV, TG and DA wrote the manuscript.

## Supporting Information

The article contains supporting Information.

**Figure S1.**
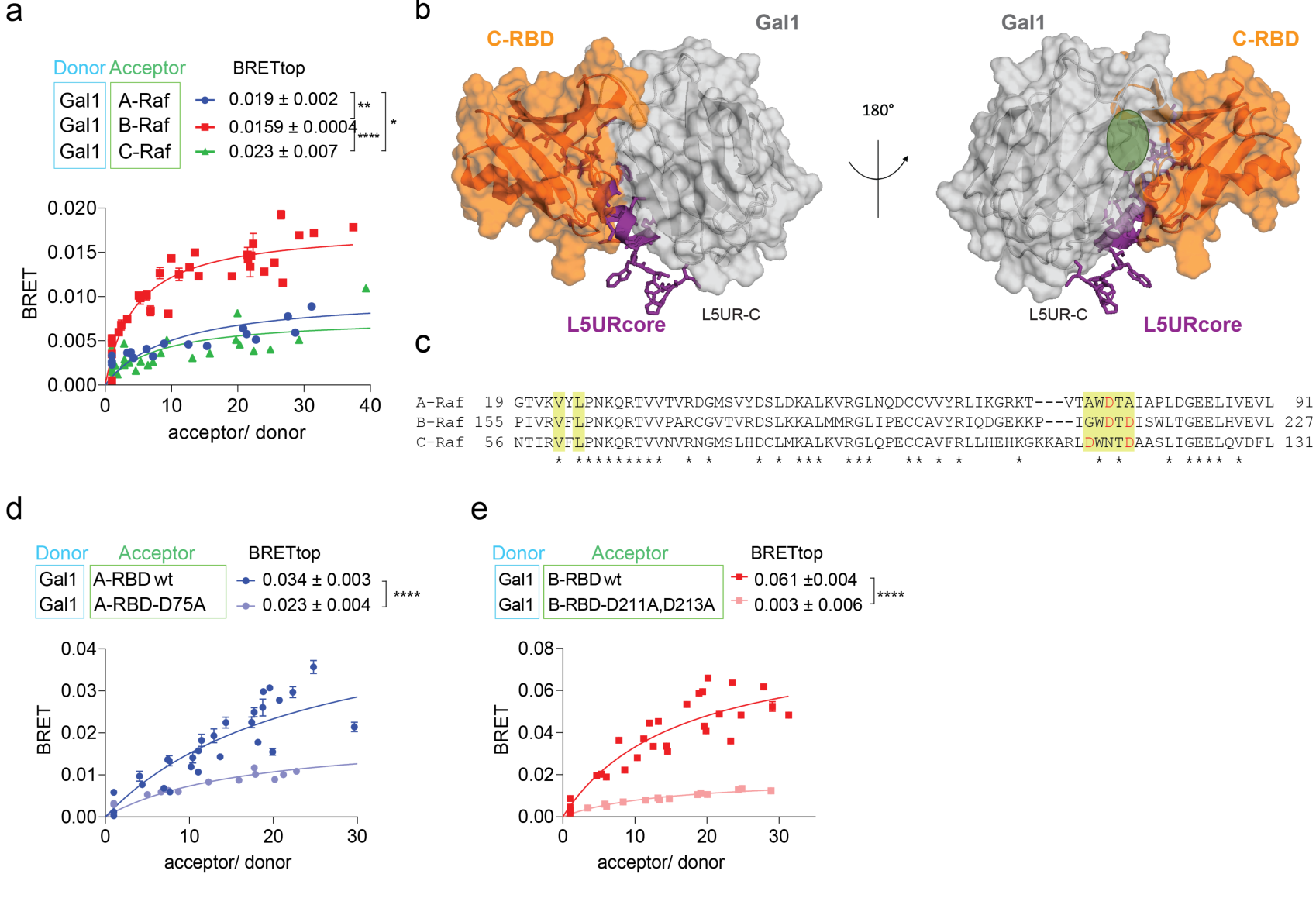
Related to main Figure 1. (**a**) BRET-titration curves of Gal1 and full-length Raf proteins; n = 3. (**b**) Computational model of hypothetical Gal1/ C-RBD/ L5UR (22-45) complex indicating the carbohydrate binding site of Gal1 (PDB ID 3W58) in green. The structural model was created with PyMOL Molecular Graphics System (Version 2.5.1) using the Gal1/ C-RBD docking model described in ^27^ and the Haddock model of Gal1/ L5UR(22-45) as described in ^34^. (**c**) Multiple sequence alignment of RBDs of A-, B- and C-Raf. The protein sequences of RBDs from the three Raf proteins, A-Raf (P10398), B-Raf (P15056) and C-Raf (P04049) were essentially as employed in the cellular assays; in brackets Uniprot database (http://uniprot.org/) accession numbers. Multiple sequence alignment was performed using Clustal Omega (https://www.ebi.ac.uk/Tools/msa/clustalo/). Yellow highlighted residues were identified as possible interaction sites with Gal1 before ^27^, and mutations tested in the BRET experiments in (d, e) are in red. (**d, e**) BRET-titration curves of Gal1 with wild-type (wt) A-RBD and A-RBD-D75A mutant (d); n = 3, or with wt B-RBD and B-RBD-D211A D213A mutant (e); n = 3.

**Figure S2.**
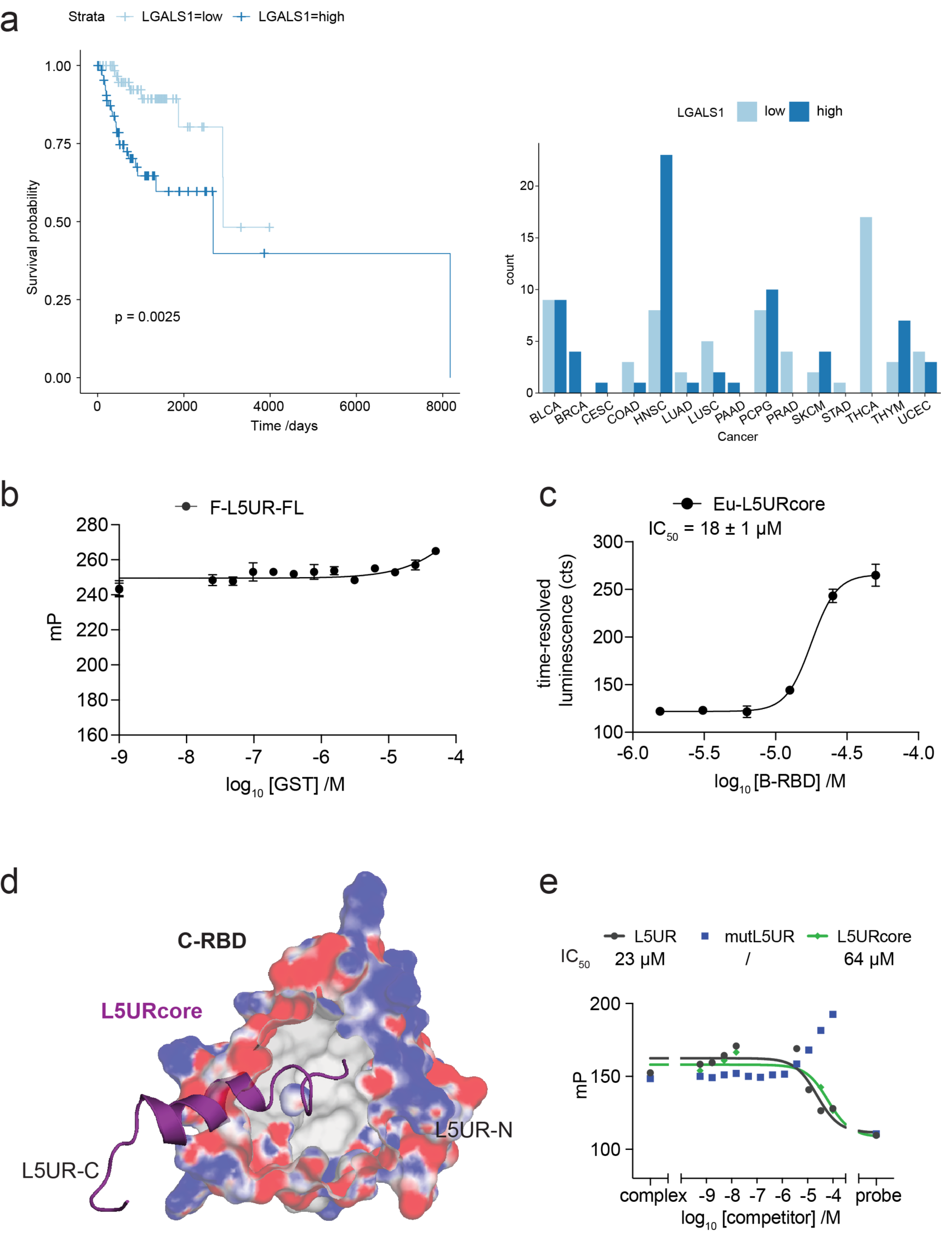
Related to main Figures 2 and 3. (**a**) PanCanAtlas data analysis reveals that high Gal1 (gene LGALS1) levels significantly decrease survival in HRAS mutant cancer cases (left). Higher Gal1 levels are more often found in head and neck (HNSC) cancers and to some extent in skin (SKCM) and thymus (THYM) cancers. These cancer types could therefore be particularly interesting for treatment with a Gal1/ Raf-interface inhibitor, which would abrogate the stimulating effect of Gal1 on oncogenic H-Ras nanoclustering and thus MAPK-signalling. (**b**) Control showing negligible binding of 10 nM F-L5UR to GST measured by fluorescence polarisation; n = 2. (**c**) Eu-L5URcore (29 nM) binding to B-RBD measured in the QRET assay using timeresolved luminescence detection. (**d**) Computational model showing putative interaction patch of the L5UR (22-45) on the CRBD. The structural model was created with PyMOL Molecular Graphics System (Version 2.5.1) using the structure of C-Raf RBD (PDB ID 1C1Y) and L5URcore (residues 22-45 of L5UR) peptide (PDB ID 2LKQ) retrieved from PDB data base (https://www.rcsb.org). We postulate a negatively charged patch (red) on the RBD at the RBD/ Gal1 interface as potential binding site for L5UR. (**e**) Displacement of F-L5UR (5 nM) from B-RBD (200 nM) by L5UR-derived peptides; n = 1.

**Figure S3.**
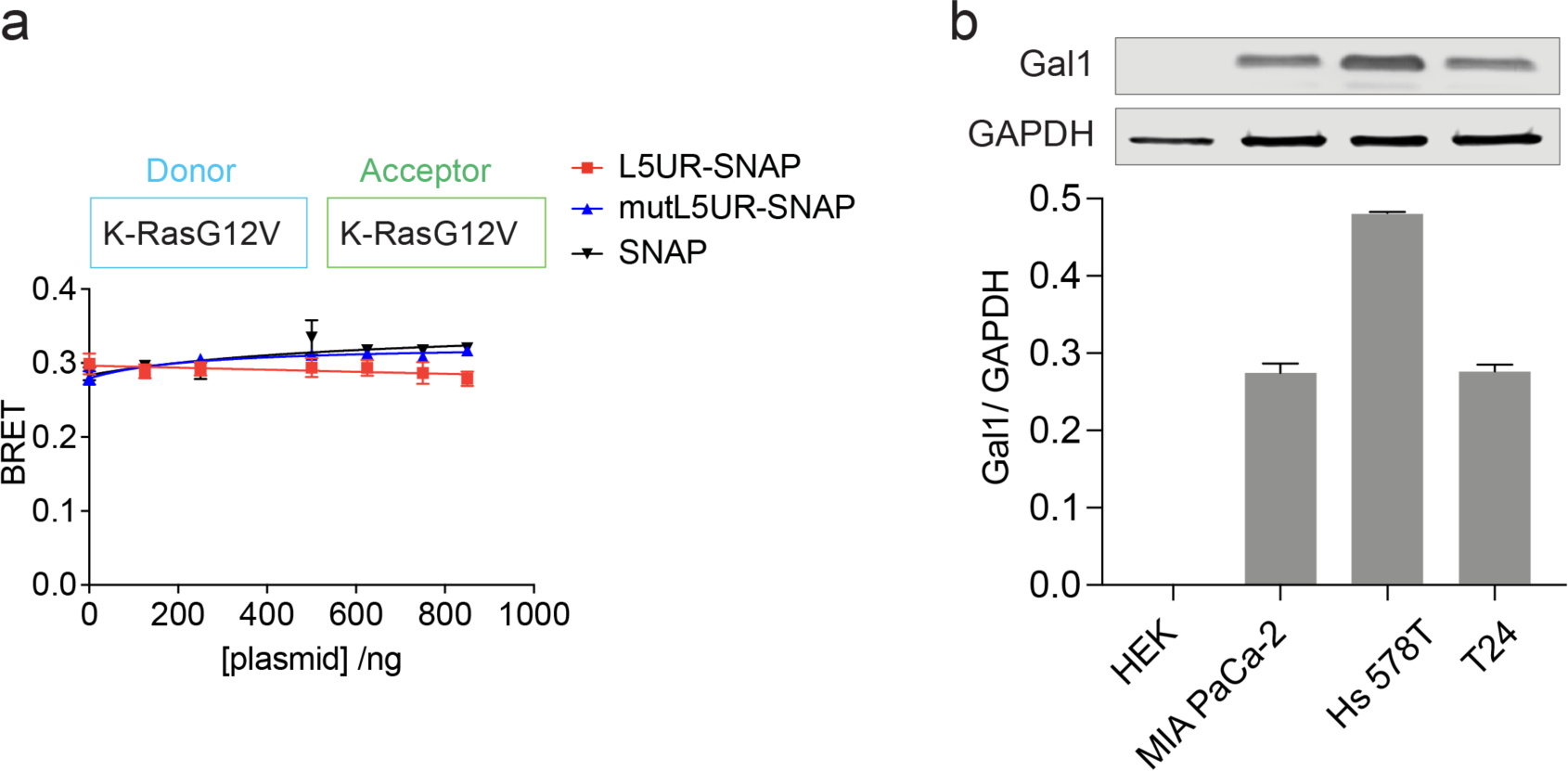
Related to main Figures 3, 4 and 5. (**a**) Negligible effect of L5UR construct expression (48 h) on K-RasG12V nanoclustering-BRET (donor:acceptor plasmid ratio = 1:10); n = 3. (**b**) Immunoblot data and quantification of endogenous Gal1 expression in employed cell lines; n = 3.

**Table S1:**
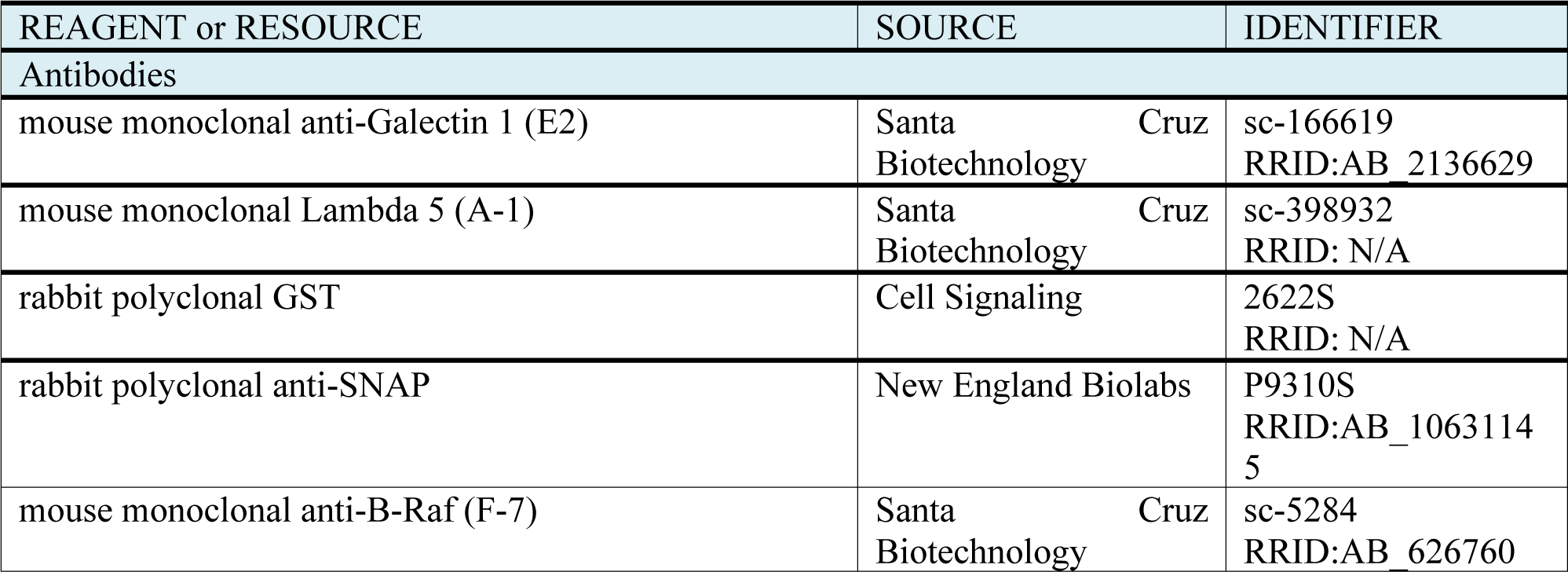
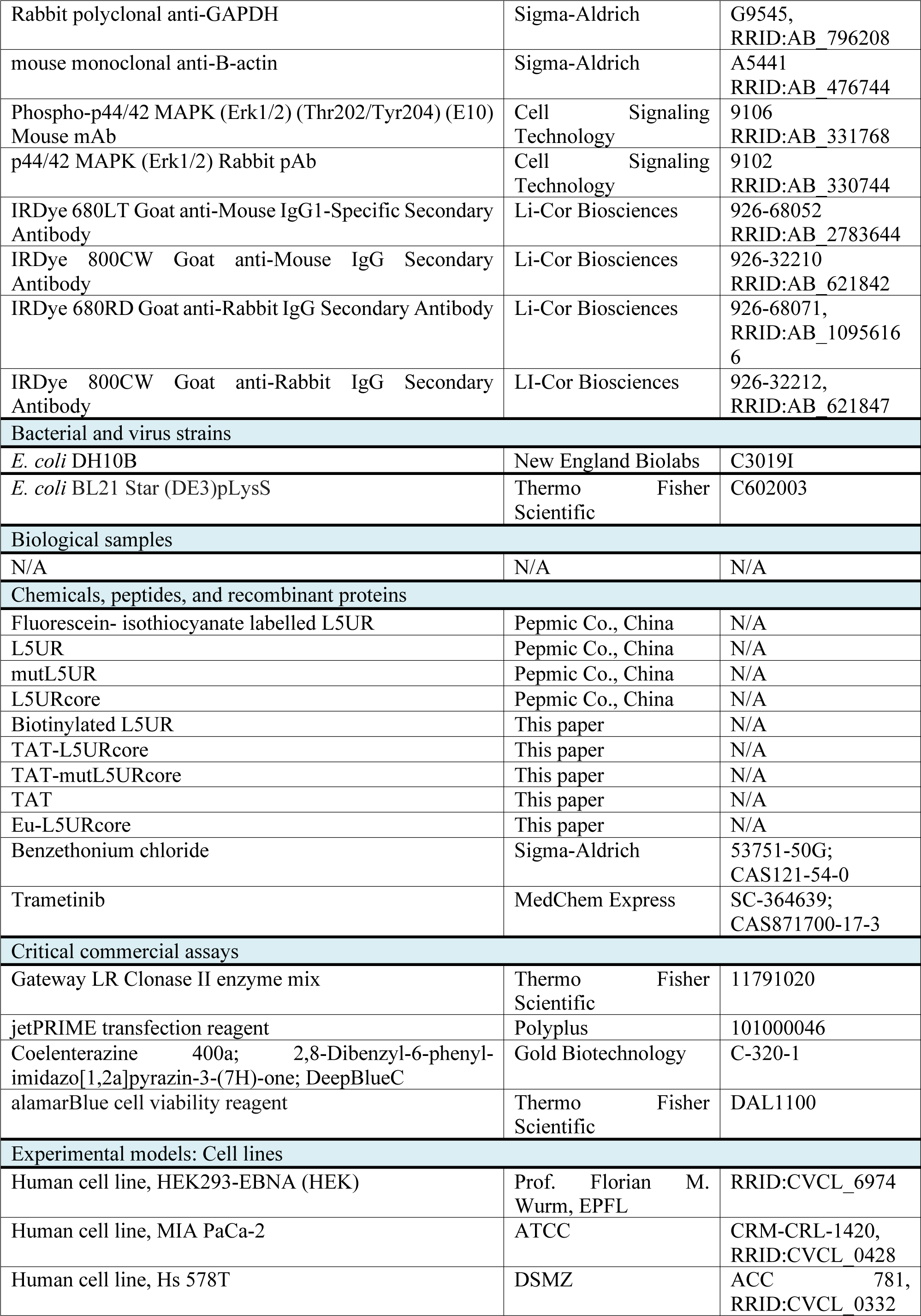
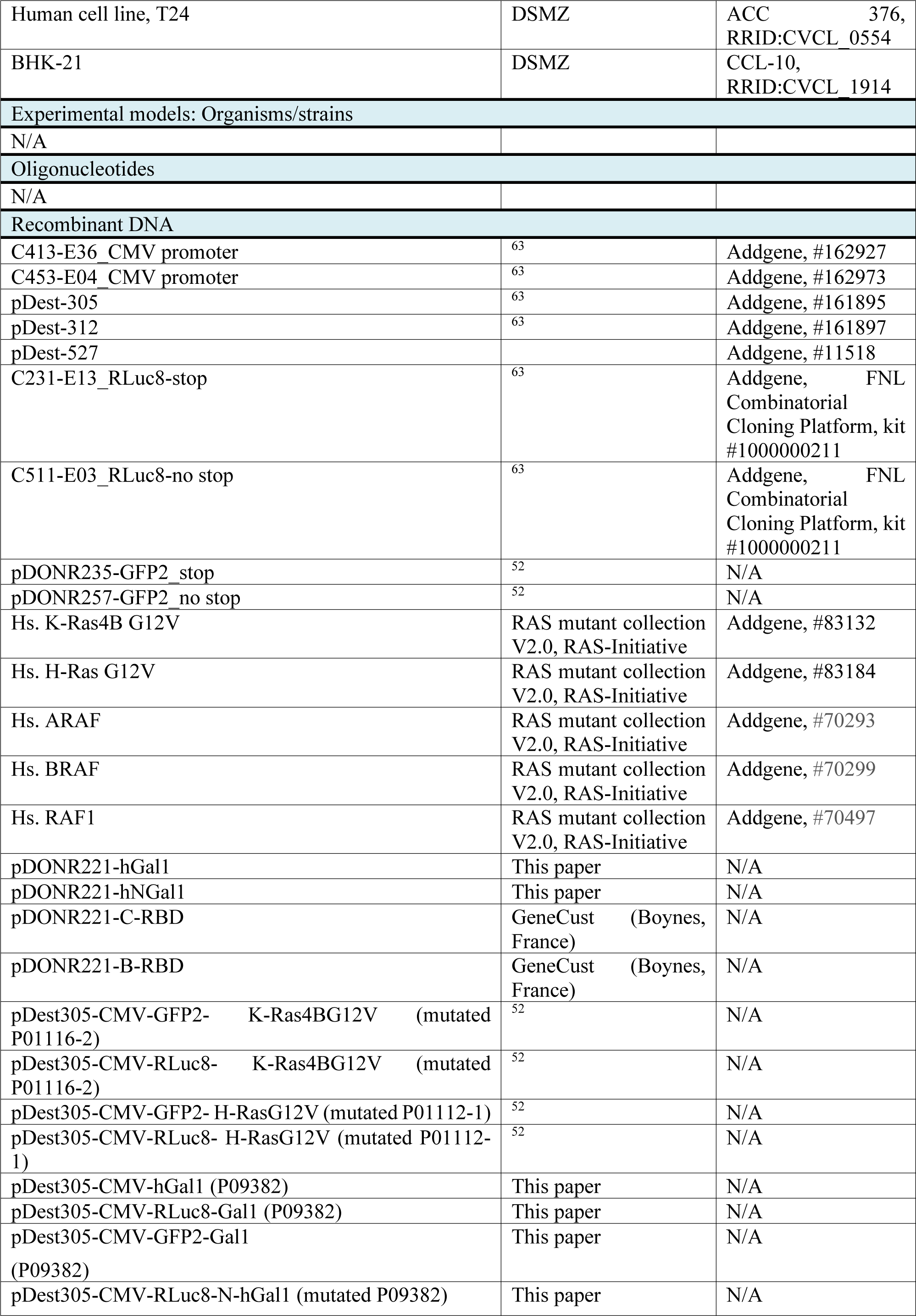
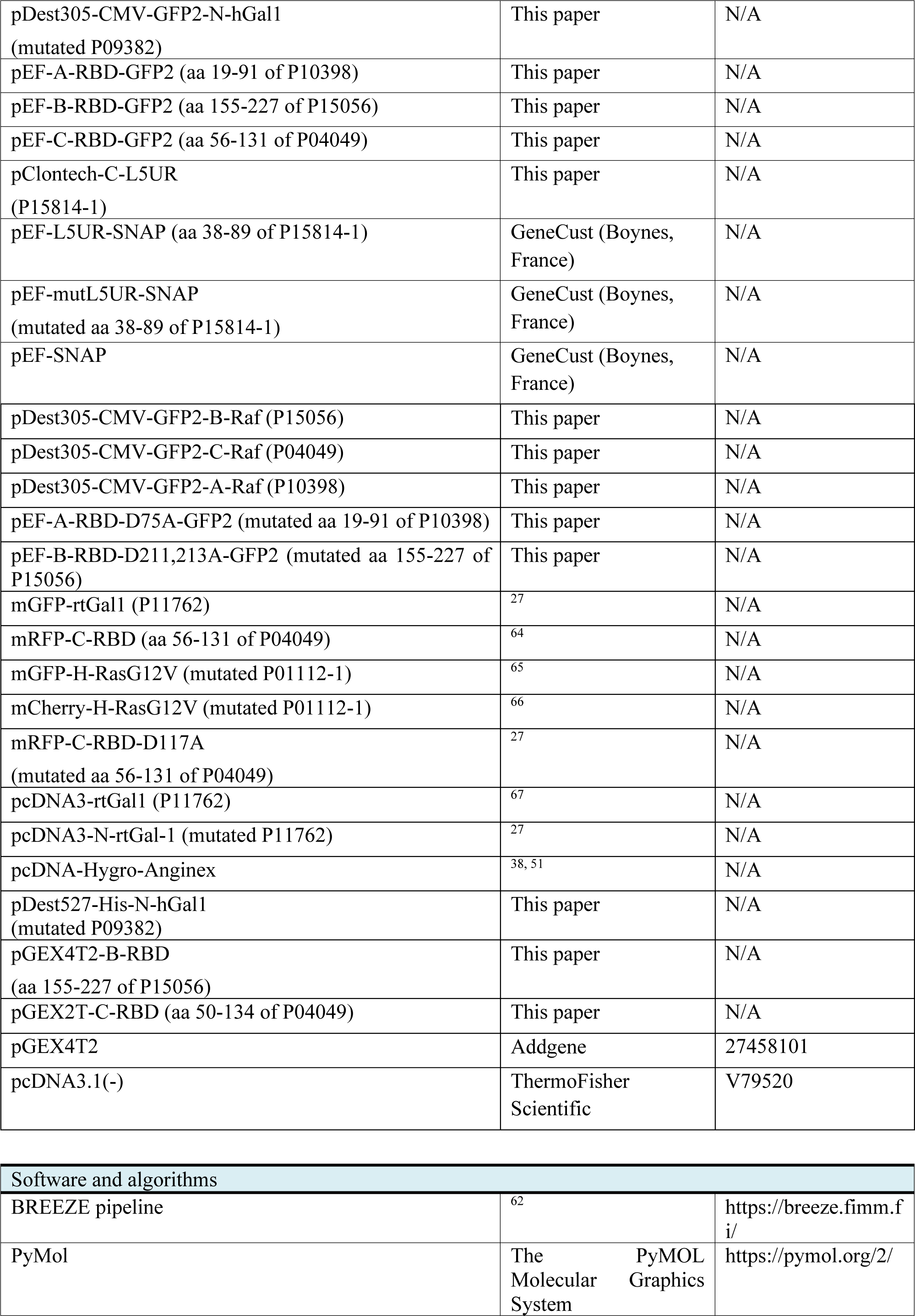
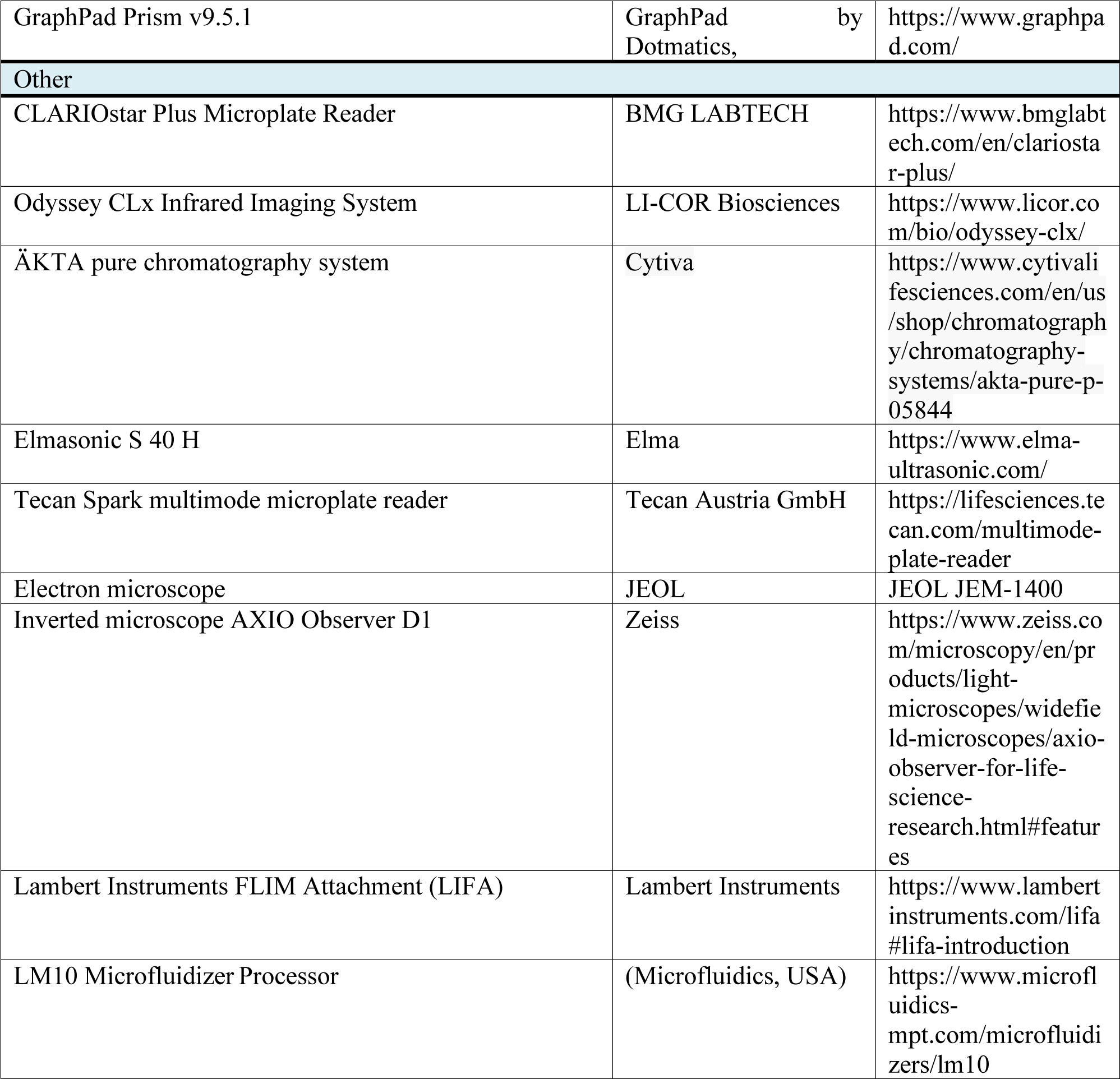
Materials and equipment employed in the study.

## References

1. Steffen CL, Kaya P, Schaffner-Reckinger E, Abankwa D. Eliminating oncogenic RAS: back to the future at the drawing board. Biochem Soc Trans 51, 447–456 (2023).

2. Punekar SR, Velcheti V, Neel BG, Wong KK. The current state of the art and future trends in RAS-targeted cancer therapies. Nat Rev Clin Oncol 19, 637–655 (2022).

3. Simanshu DK, Nissley DV, McCormick F. RAS Proteins and Their Regulators in Human Disease. Cell 170, 17–33 (2017).

4. Spiegel J, Cromm PM, Zimmermann G, Grossmann TN, Waldmann H. Small-molecule modulation of Ras signaling. Nat Chem Biol 10, 613–622 (2014).

5. Simanshu DK, Morrison DK. A Structure is Worth a Thousand Words: New Insights for RAS and RAF Regulation. Cancer Discov 12, 899–912 (2022).

6. Lavoie H, Therrien M. Regulation of RAF protein kinases in ERK signalling. Nat Rev Mol Cell Biol 16, 281–298 (2015).

7. Martinez Fiesco JA, Durrant DE, Morrison DK, Zhang P. Structural insights into the BRAF monomer-to-dimer transition mediated by RAS binding. Nat Commun 13, 486 (2022).

8. Rajakulendran T, Sahmi M, Lefrancois M, Sicheri F, Therrien M. A dimerization-dependent mechanism drives RAF catalytic activation. Nature 461, 542–545 (2009).

9. Abankwa D, Gorfe AA. Mechanisms of Ras Membrane Organization and Signaling: Ras Rocks Again. Biomolecules 10, (2020).

10. Abankwa D, Gorfe AA, Hancock JF. Ras nanoclusters: molecular structure and assembly. Semin Cell Dev Biol 18, 599–607 (2007).

11. Plowman SJ, Hancock JF. Ras signaling from plasma membrane and endomembrane microdomains. Biochim Biophys Acta 1746, 274–283 (2005).

12. Tian T, Harding A, Inder K, Plowman S, Parton RG, Hancock JF. Plasma membrane nanoswitches generate high-fidelity Ras signal transduction. Nat Cell Biol 9, 905–914 (2007).

13. Sarkar-Banerjee S, Sayyed-Ahmad A, Prakash P, Cho KJ, Waxham MN, Hancock JF, Gorfe AA. Spatiotemporal Analysis of K-Ras Plasma Membrane Interactions Reveals Multiple High Order Homo-oligomeric Complexes. J Am Chem Soc 139, 13466–13475 (2017).

14. Cho KJ, et al. Raf inhibitors target ras spatiotemporal dynamics. Curr Biol 22, 945–955 (2012).

15. Jin T, Lavoie H, Sahmi M, David M, Hilt C, Hammell A, Therrien M. RAF inhibitors promote RAS-RAF interaction by allosterically disrupting RAF autoinhibition. Nat Commun 8, 1211 (2017).

16. Holderfield M, Deuker MM, McCormick F, McMahon M. Targeting RAF kinases for cancer therapy: BRAF-mutated melanoma and beyond. Nat Rev Cancer 14, 455–467 (2014).

17. Pavic K, Chippalkatti R, Abankwa D. Drug targeting opportunities en route to Ras nanoclusters. Adv Cancer Res 153, 63–99 (2022).

18. Rotblat B, et al. H-Ras nanocluster stability regulates the magnitude of MAPK signal output. PLoS One 5, e11991 (2010).

19. Prior IA, Muncke C, Parton RG, Hancock JF. Direct visualization of Ras proteins in spatially distinct cell surface microdomains. J Cell Biol 160, 165–170 (2003).

20. Elad-Sfadia G, Haklai R, Ballan E, Gabius HJ, Kloog Y. Galectin-1 augments Ras activation and diverts Ras signals to Raf-1 at the expense of phosphoinositide 3-kinase. J Biol Chem 277, 37169–37175 (2002).

21. Timoshenko AV. Towards molecular mechanisms regulating the expression of galectins in cancer cells under microenvironmental stress conditions. Cell Mol Life Sci 72, 4327–4340 (2015).

22. Rabinovich GA. Galectin-1 as a potential cancer target. Br J Cancer 92, 1188–1192 (2005).

23. Johannes L, Jacob R, Leffler H. Galectins at a glance. J Cell Sci 131, (2018).

24. Belanis L, Plowman SJ, Rotblat B, Hancock JF, Kloog Y. Galectin-1 is a novel structural component and a major regulator of h-ras nanoclusters. Mol Biol Cell 19, 1404–1414 (2008).

25. Mejuch T, van Hattum H, Triola G, Jaiswal M, Waldmann H. Specificity of Lipoprotein Chaperones for the Characteristic Lipidated Structural Motifs of their Cognate Lipoproteins. Chembiochem 16, 2460–2465 (2015).

26. Lakshman B, et al. Quantitative biophysical analysis defines key components modulating recruitment of the GTPase KRAS to the plasma membrane. J Biol Chem 294, 2193–2207 (2019).

27. Blazevits O, et al. Galectin-1 dimers can scaffold Raf-effectors to increase H-ras nanoclustering. Sci Rep 6, 24165 (2016).

28. Siljamaki E, Abankwa D. SPRED1 Interferes with K-ras but Not H-ras Membrane Anchorage and Signaling. Mol Cell Biol 36, 2612–2625 (2016).

29. Stegmayr J, et al. Extracellular and intracellular small-molecule galectin-3 inhibitors. Sci Rep 9, 2186 (2019).

30. Chan YC, Lin HY, Tu Z, Kuo YH, Hsu SD, Lin CH. Dissecting the Structure-Activity Relationship of Galectin-Ligand Interactions. Int J Mol Sci 19, (2018).

31. Marullo S, Bouvier M. Resonance energy transfer approaches in molecular pharmacology and beyond. Trends Pharmacol Sci 28, 362–365 (2007).

32. Manoharan GB, Laurini C, Bottone S, Ben Fredj N, Abankwa DK. K-Ras Binds Calmodulin-Related Centrin1 with Potential Implications for K-Ras Driven Cancer Cell Stemness. Cancers (Basel) 15, (2023).

33. Cho M, Cummings RD. Galectin-1, a beta-galactoside-binding lectin in Chinese hamster ovary cells. I. Physical and chemical characterization. J Biol Chem 270, 5198–5206 (1995).

34. Elantak L, et al. Structural basis for galectin-1-dependent pre-B cell receptor (pre-BCR) activation. J Biol Chem 287, 44703–44713 (2012).

35. Dings RP, et al. Antitumor agent calixarene 0118 targets human galectin-1 as an allosteric inhibitor of carbohydrate binding. J Med Chem 55, 5121–5129 (2012).

36. Astorgues-Xerri L, et al. OTX008, a selective small-molecule inhibitor of galectin-1, downregulates cancer cell proliferation, invasion and tumour angiogenesis. Eur J Cancer 50, 2463–2477 (2014).

37. Brandwijk RJ, Nesmelova I, Dings RP, Mayo KH, Thijssen VL, Griffioen AW. Cloning an artificial gene encoding angiostatic anginex: From designed peptide to functional recombinant protein. Biochem Biophys Res Commun 333, 1261–1268 (2005).

38. Thijssen VL, et al. Galectin-1 is essential in tumor angiogenesis and is a target for antiangiogenesis therapy. Proc Natl Acad Sci U S A 103, 15975–15980 (2006).

39. Jauset T, Beaulieu ME. Bioactive cell penetrating peptides and proteins in cancer: a bright future ahead. Curr Opin Pharmacol 47, 133–140 (2019).

40. Dietrich L, et al. Cell Permeable Stapled Peptide Inhibitor of Wnt Signaling that Targets beta-Catenin Protein-Protein Interactions. Cell Chem Biol 24, 958–968 e955 (2017).

41. Vives E, Brodin P, Lebleu B. A truncated HIV-1 Tat protein basic domain rapidly translocates through the plasma membrane and accumulates in the cell nucleus. J Biol Chem 272, 16010–16017 (1997).

42. Adihou H, et al. A protein tertiary structure mimetic modulator of the Hippo signalling pathway. Nat Commun 11, 5425 (2020).

43. Siddiqui FA, Vukic V, Salminen TA, Abankwa D. Elaiophylin Is a Potent Hsp90/ Cdc37 Protein Interface Inhibitor with K-Ras Nanocluster Selectivity. Biomolecules 11, (2021).

44. Siddiqui FA, et al. Novel Small Molecule Hsp90/Cdc37 Interface Inhibitors Indirectly Target K-Ras-Signaling. Cancers (Basel) 13, (2021).

45. Shalom-Feuerstein R, Plowman SJ, Rotblat B, Ariotti N, Tian T, Hancock JF, Kloog Y. K-ras nanoclustering is subverted by overexpression of the scaffold protein galectin-3. Cancer Res 68, 6608–6616 (2008).

46. Terrell EM, et al. Distinct Binding Preferences between Ras and Raf Family Members and the Impact on Oncogenic Ras Signaling. Mol Cell 76, 872–884 e875 (2019).

47. Spencer-Smith R, et al. Inhibition of RAS function through targeting an allosteric regulatory site. Nat Chem Biol 13, 62–68 (2017).

48. Prior IA, Hood FE, Hartley JL. The Frequency of Ras Mutations in Cancer. Cancer Res 80, 2969–2974 (2020).

49. Burtness B, et al. Pembrolizumab alone or with chemotherapy versus cetuximab with chemotherapy for recurrent or metastatic squamous cell carcinoma of the head and neck (KEYNOTE-048): a randomised, open-label, phase 3 study. Lancet 394, 1915–1928 (2019).

50. Ho AL, et al. Tipifarnib in Head and Neck Squamous Cell Carcinoma With HRAS Mutations. J Clin Oncol 39, 1856–1864 (2021).

51. Brandwijk RJ, Dings RP, van der Linden E, Mayo KH, Thijssen VL, Griffioen AW. Anti-angiogenesis and anti-tumor activity of recombinant anginex. Biochem Biophys Res Commun 349, 1073–1078 (2006).

52. Okutachi S, et al. A Covalent Calmodulin Inhibitor as a Tool to Study Cellular Mechanisms of K-Ras-Driven Stemness. Front Cell Dev Biol 9, 665673 (2021).

53. Manoharan GB, Okutachi S, Abankwa D. Potential of phenothiazines to synergistically block calmodulin and reactivate PP2A in cancer cells. PLoS One 17, e0268635 (2022).

54. Manoharan GB, Kopra K, Eskonen V, Harma H, Abankwa D. High-throughput amenable fluorescence-assays to screen for calmodulin-inhibitors. Anal Biochem 572, 25–32 (2019).

55. Sinijarv H, Wu S, Ivan T, Laasfeld T, Viht K, Uri A. Binding assay for characterization of protein kinase inhibitors possessing sub-picomolar to sub-millimolar affinity. Anal Biochem 531, 67–77 (2017).

56. Harma H, et al. A new simple cell-based homogeneous time-resolved fluorescence QRET technique for receptor-ligand interaction screening. J Biomol Screen 14, 936–943 (2009).

57. Kopra K, Harma H. Quenching resonance energy transfer (QRET): a single-label technique for inhibitor screening and interaction studies. N Biotechnol 32, 575–580 (2015).

58. Kopra K, et al. A homogeneous quenching resonance energy transfer assay for the kinetic analysis of the GTPase nucleotide exchange reaction. Anal Bioanal Chem 406, 4147–4156 (2014).

59. Guzman C, Oetken-Lindholm C, Abankwa D. Automated High-Throughput Fluorescence Lifetime Imaging Microscopy to Detect Protein-Protein Interactions. J Lab Autom 21, 238–245 (2016).

60. Guzman C, et al. The efficacy of Raf kinase recruitment to the GTPase H-ras depends on H-ras membrane conformer-specific nanoclustering. J Biol Chem 289, 9519–9533 (2014).

61. Babu Manoharan G, Guzman C, Najumudeen AK, Abankwa D. Detection of Ras nanoclustering-dependent homo-FRET using fluorescence anisotropy measurements. Eur J Cell Biol 102, 151314 (2023).

62. Potdar S, et al. Breeze 2.0: an interactive web-tool for visual analysis and comparison of drug response data. Nucleic Acids Res, (2023).

63. Wall VE, Garvey LA, Mehalko JL, Procter LV, Esposito D. Combinatorial assembly of clone libraries using site-specific recombination. Methods Mol Biol 1116, 193–208 (2014).

64. Abankwa D, Gorfe AA, Inder K, Hancock JF. Ras membrane orientation and nanodomain localization generate isoform diversity. Proc Natl Acad Sci U S A 107, 1130–1135 (2010).

65. Abankwa D, et al. A novel switch region regulates H-ras membrane orientation and signal output. EMBO J 27, 727–735 (2008).

66. Solman M, et al. Specific cancer-associated mutations in the switch III region of Ras increase tumorigenicity by nanocluster augmentation. Elife 4, e08905 (2015).

67. Paz A, Haklai R, Elad-Sfadia G, Ballan E, Kloog Y. Galectin-1 binds oncogenic H-Ras to mediate Ras membrane anchorage and cell transformation. Oncogene 20, 7486–7493 (2001).

